# Programming cell behavior with synthetic protease-activated receptors

**DOI:** 10.64898/2026.06.07.730697

**Authors:** Matthew Ravalin, Nicholas A. Kalogriopoulos, Rocco Latorre, Gabriele Kockelkoren, Reika Tei, Martina Chieca, Francesco De Logu, Nigel W Bunnett, Alice Y. Ting

**Affiliations:** Department of Genetics, Stanford University, Stanford, CA, USA; Department of Molecular Pathobiology, College of Dentistry, New York University, New York, NY, United States; Pain Research Center, College of Dentistry, New York University, New York, NY, United States; Department of Health Sciences, Clinical Pharmacology and Oncology Section, University of Florence, Florence, 50139, Italy; Department of Biology, Stanford University, Stanford, CA, USA; Department of Chemistry, Stanford University, Stanford, CA, USA; Chan Zuckerberg Biohub—San Francisco, San Francisco, CA, USA; Phil & Penny Knight Initiative for Brain Resilience at the Wu Tsai Neurosciences Institute, Stanford University, Stanford, CA, USA

## Abstract

Extracellular proteases are important signaling molecules in coagulation, inflammation, cell migration, and pain. Dysregulation of extracellular protease activity is common in diseases that perturb these critical functions. Engineering cells to sense and respond programmatically to protease activity has applications in biosensing, cell-based screening for protease activity, and therapeutics. Here we report synthetic protease-activated receptors (**SynPARs**) based on engineered, auto-inhibited G protein-coupled receptors (GPCRs). Relief of autoinhibition by proteolysis enables receptor activation by an exogenous or tethered agonist to generate transgene expression, real-time fluorescence, or endogenous G-protein signaling. We demonstrate SynPAR modularity with diverse secreted proteases, establish a cell-based SynPAR library selection to optimize protease recognition sequences, and control neuronal activity in response to protease activity. Finally, we use SynPARs in the dorsal root ganglion of mice to counteract hyperalgesia produced by trypsin activity, rewiring neurons to produce an analgesic response to a pain-inducing stimulus. Our study establishes SynPAR as a versatile and modular platform for recording, sensing, and responding to pericellular proteolysis. This fills a critical gap in protease-sensing tools and lays the groundwork for protease-activated genetic and cell-based medicines.

## Introduction

Nearly half of the approximately 600 human proteases are secreted into the extracellular milieu where they function in autocrine, paracrine, and endocrine signaling.^1^ Proteolysis is an irreversible post-translational modification. As such, proteases are highly regulated by localization, pH, production as inactive zymogens, and an ensemble of endogenous protease inhibitors. These and more exotic layers of regulation work in concert to restrict protease activity to the appropriate time and place for signaling. The decay of these regulatory mechanisms contributes to disease states including coagulopathies, inflammatory conditions, tumor metastasis, and chronic pain. The intricate and distributed regulatory mechanisms that control extracellular protease signaling make modeling protease biology *in vitro* a significant challenge. Moreover, the central role of enzyme activity rather than abundance makes sequencing and proteomic methods insufficient to capture the dynamics of protease biology. Activity-based probes and biosensors have provided insight into the spatiotemporal regulation of proteases, albeit with a limited scope of functional outputs. ^2,3^ Synthetic receptors enable a range of recording, sensing, and response functionalities inaccessible to other methods.^4^ However, existing platforms measure the abundance of proteins and lack the capacity to interrogate enzyme activity. To address this technological gap, we developed a modular synthetic receptor platform that can detect the activity of an extracellular protease and convert that activity into an array of functional outputs (**Figure 1A**).

**Figure 1.**
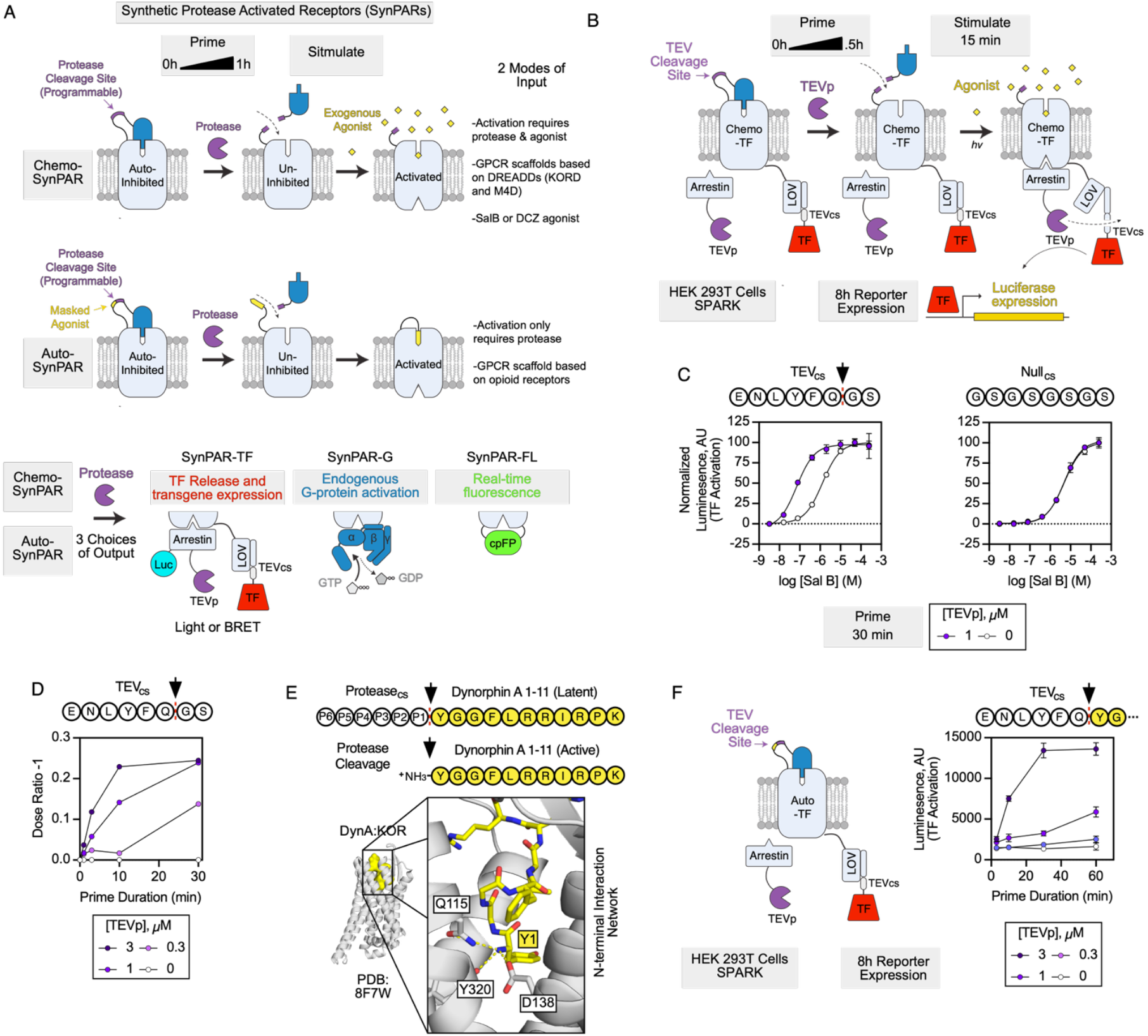
SynPARs integrate pericellular protease activity to drive defined outputs. **(A)** Synthetic protease activated receptors (SynPARs) are comprised of a protease-cleavable autoinhibitory domain tethered to a GPCR. Cleavage of the input module relieves inhibition, allowing either an exogenous agonist (Chemo-SynPAR, top) or a masked peptide agonist (Auto-SynPAR, middle) to activate the GPCR. This conditional activity can be coupled to a variety of outputs (bottom) including light/BRET-gated transgene expression (SynPAR_TF_), endogenous G-protein coupling (SynPAR_G_), and real-time fluorescence (SynPAR_FL_).Luc, luciferase. TEVp, TEV protease. TF, transcription factor. cpFP, circularly permuted fluorescent protein. **(B)** Detail for TEV_cs_-Chemo-SynPAR_TF_, which couples extracellular TEV protease activity to transgene expression based on SPARK (specific protein association tool giving transcriptional readout with rapid kinetics.^8^ TEVcs, TEVp cleavage site. Extracellular SynPAR cleavage (Prime step) followed by coincident light and agonist (Stimulate step) leads to intracellular proteolytic release of a tethered transcription factor and reporter gene (e.g., firefly luciferase) expression. **(C)** Treatment of TEV_cs_-Chemo-SynPAR_TF_ with 1 μM TEVp for 30 min sensitizes the receptor to agonist (SalB). Identical treatment of a construct that has a Gly-Ser linker instead of a cleavage site (Null_cs_) does not change the response to agonist. Data plotted as mean of triplicate, with error plotted as ±1 std. dev. This experiment was performed 2 times. **(D)** Sensitization can be compared across conditions using the Dose Ratio-1 (pEC50_(+)PROTEASE_/pEC50_(-)PROTEASE_ -1). Raw data in **Figure S2B**. This experiment was performed 2 times. **(E)** A protease cleavage site (P1-P6) masks the N-terminus of the opioid peptide Dynorphin A (DynA _1-11_, yellow), which binds the *k*-opioid receptor (*k*OR) in its orthosteric site (Q115, Y320, and D138) via a hydrogen bonding network with Dynorphin’s N-terminal amine. **(F)** TEV_cs_-Auto-SynPAR_TF_ (Left) exhibits saturable time- and concentration-dependent response to TEV protease activity (Right). Luminescence plotted as a mean of triplicates with error expressed as ±1 std. dev. This experiment was performed 2 times.

Among natural G-protein coupled receptors (GPCRs), protease-activated receptors (PARs, **Figure S1**) are critical components of signaling networks that control platelet aggregation, inflammation, and nociception. PARs are activated when a protease cleaves the extracellular N-terminus of the receptor, unmasking a latent agonist that binds in *cis* and activates intracellular G-protein coupled signaling cascades.^5^ Inspired by PARs and our recent work developing programmable antigen-gated engineered G-protein coupled receptors (**PAGERs, Figure S2**), we developed SynPARs (Synthetic Protease Activated Receptors) to convert a defined endoproteolytic activity into programmable functional outputs.^6^ SynPARs can function under chemogenetic (**Chemo-SynPAR**) or autonomous (**Auto-SynPAR**) control to generate transgene expression (**SynPAR**_**TF**_), real-time fluorescence (**SynPAR**_**FL**_), or G-protein activation (**SynPAR**_**G**_). Collectively, SynPARs comprise a powerful platform for recording, sensing, and responding to pericellular protease activity.

### SynPARs integrate extracellular protease activity

We sought to emulate and expand on the activation logic of PARs in a system that is amenable to reprogramming the input and output activities. We developed two complementary SynPAR platforms. In the first (Chemo-SynPAR, **Figure 1A, Top**), chemogenetic control is imparted by leveraging a mechanism that depends on the relief of autoinhibition. We generated a protease-labile autoinhibitory module that consists of an N-terminal antagonist, a protease cleavage site, and a tether to connect the antagonist to the GPCR of choice. Cleavage leads to dissociation of the antagonist and sensitization to exogenous agonist. In the second (Auto-SynPAR, **Figure 1A, Middle**), we achieve autonomous activation by coupling cleavage to the concerted relief of autoinhibition and the generation of a tethered agonist. Proper pairing of antagonist, agonist, inhibitory modules and a GPCR enable Chemo and Auto-SynPARs to access multiple output mechanisms (**Figure 1A, Bottom**).

To generate a prototype Chemo-SynPAR, we tethered an inhibitor of the kappa opioid receptor DREADD (KORD) to its N-terminus and incorporated a tobacco etch virus protease cleavage sequence (TEV_cs_, ENLYFQ/GS) in the linker.^7^ We coupled receptor activation to the release of transcription factor (TF) using SPARK (**Figure 1B**).^8^ To test the sensitivity and specificity of TEV_cs_-Chemo-SynPAR_TF_, we compared the EC_50_ of salvinorin B (SalB, KORD agonist) in cells treated with 1 μM recombinant TEV_p_ for 30 min (Priming) to cells not treated with TEV_p_. In parallel, we treated cells expressing a similar construct that contained a Gly-Ser linker instead of TEV_cs_ (Null_cs_-Chemo-SynPAR_TF_). After exposure to SalB and Light (Stimulation) and incubation for 8h, cells expressing TEV_cs_-Chemo-SynPAR_TF_ and treated with TEV_p_ exhibited a leftward shift in SalB EC_50_. Cells expressing Null_cs_-Chemo-SynPAR_TF_ showed no such shift (**Figure 1C**). To quantitatively compare the relative activation of these constructs we calculated the dose ratio (DR=pEC_50-Treated_/pEC_50-Untreated_) for each construct (**Figure S2A**). With specificity established, we varied the protease treatment time and concentration to determine how SynPAR responds to different levels of protease activity. These data demonstrated that Chemo-SynPAR_TF_ integrates protease activity in a time and dose dependent manner (**Figure 1D, Figure S2B, C**) and that fractional cleavage of the receptor and the reporter activation are closely linked (**Figure S2D**).

Chemogenetic tools afford temporal control.^9^ However, the delivery of a chemogenetic trigger also complicates the use of these systems *in vivo*, particularly in therapeutic or real-time sensing use cases. We developed Auto-SynPAR to address this liability by functioning without an exogenous agonist. Like native PARs, kappa opioid receptor (*κ*OR) peptide agonists bind to the orthosteric site via a salt bridge formed by the N-terminal amine. This interaction is mediated by Asp138 of KOR.^10^ To generate a prototype Auto-SynPAR_TF_, we first reverted the D138N mutation in the KORD component of TEV_cs_-Chemo-SynPAR_TF_ to allow activation by peptide agonists. We then nested the KOR peptide agonist dynorphin_1-11_ behind the TEV_cs_ such that TEV_p_ cleavage would yield the free N-terminal tyrosine of the agonist (**Figure 1E**). Using SPARK, we demonstrated that TEV_cs_-Auto-SynPAR_TF_ can also yield time- and concentration-dependent activity after priming with TEV_p_ and stimulation with light (**Figure 1F**). Chemo and Auto-SynPAR_TF_ both exhibit robust activation-dependent internalization, consistent with canonical GPCR behavior (**Figure S3A, B**). To establish the temporal resolution of Chemo and Auto-SynPAR_TF_, we measured the decay of response to protease treatment relative to stimulation (Light and 1 μM SalB for Chemo-SynPAR_TF_, Light for Auto-SynPAR_TF_). This demonstrated that the half-lives of Chemo-SynPAR_TF_ and Auto-SynPAR_TF_ were ∼4h and ∼2h respectively. (**Figure S4A, B**). This data suggests that the Chemo-SynPAR platform may be better suited for long-term recording of protease activities, while the Auto-SynPAR platform may be preferred for acute response to proteolytic stimuli. Together these systems provide flexible programming of cellular responses to proteolysis.

### SynPARs are a modular input-output system

Secreted proteases vary widely in their preferred substrates, catalytic efficiency, and mechanism. TEV_p_ is a bioorthogonal cysteine protease derived from a plant virus and widely used in synthetic biology.^11^ To establish the input modularity of the SynPAR platform using native human proteases, we replaced TEV_cs_ with published cleavage sequences for human thrombin, uPA, MMP9, and renin in the Chemo-SynPAR_TF_ construct (**Figure 2A**).^12^ We used NanoLuc-based BRET rather than light to uncage the LOV domain of SynPAR-TF to reduce background and maximize sensitivity.^13^ In this context we were able to see protease-dependent responses for all constructs after treatment with protease for 1h and stimulation with Furimazine (substrate for NanoLuc) and SalB for 15min (**Figure 2B**). The diversity of dynamic ranges among these constructs generated with published cleavage sequences suggests that cleavage site optimization in the context of SynPAR would be useful for targeting individual proteases of interest. For example, the MMP9 cleavage sequence reduces basal inhibition of SynPAR, while uPA-directed SynPAR shows modest turn-on and low signal in response to protease activation.

**Figure 2.**
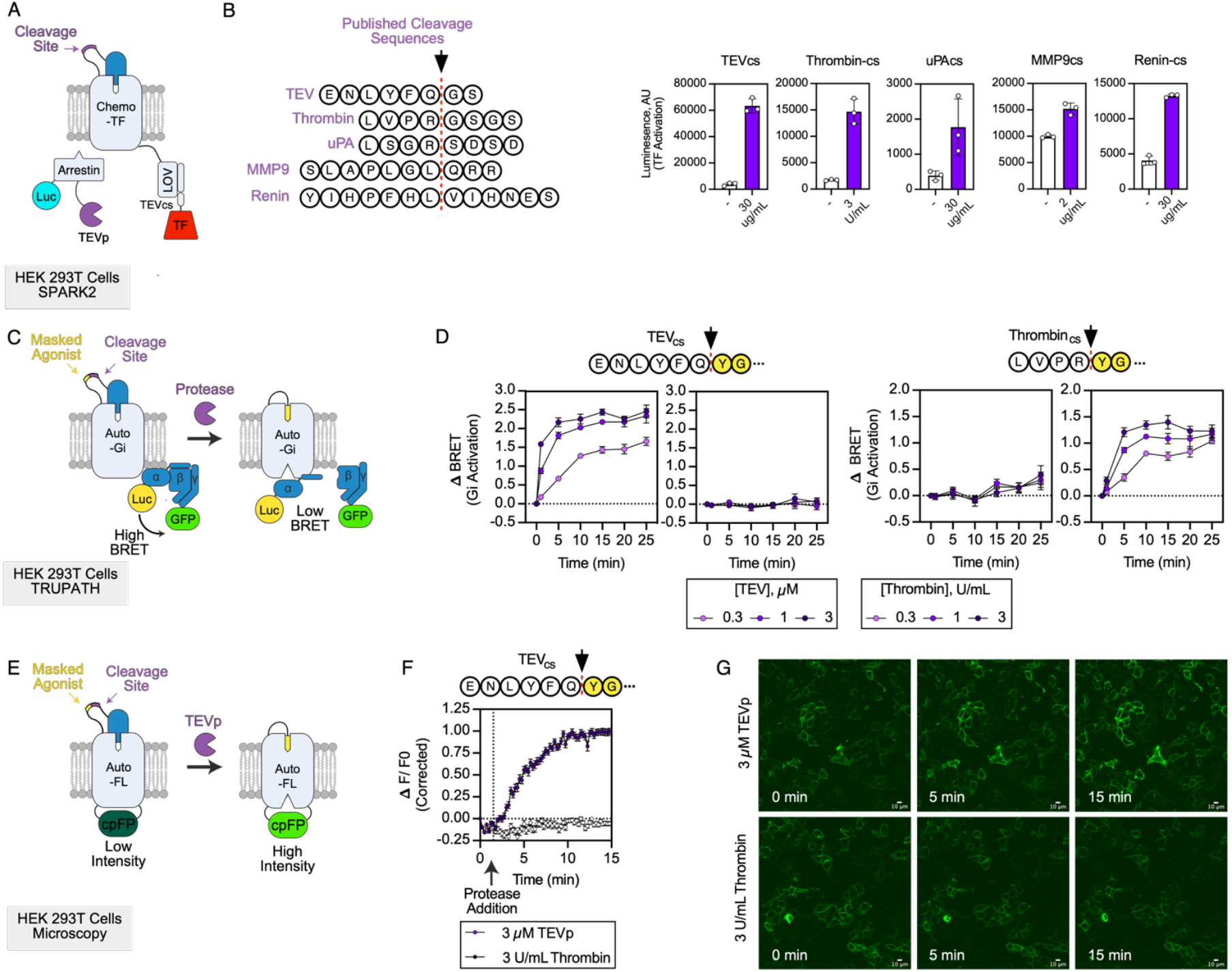
SynPAR inputs and outputs can be reprogrammed. **(A)** Schematic of Chemo-SynPAR-TF showing that the protease specificity can be reprogrammed by changing the primary amino acid sequence in the linker of the autoinhibitory domain to published substrates for target proteases of interest and read out by transgene expression based on SPARK2, which uses BRET rather than light to uncage the transcription factor.^13^ **(B)** Left: Different protease recognition sequences tested in Chemo-SynPAR-TF. Right: HEK 293T cells transiently expressing the indicated Chemo-SynPAR-TF, NanoLuc-arrestin-TEVp, and UAS-FLuc were treated with the corresponding protease, SalB, and furimazine (to initiate BRET from NanoLuc to LOV-TEVcs). Right: Data plotted as mean of triplicate (bar), with individual data points shown. Errors, ±1 std. dev. This experiment was performed 2 times **(C)** Schematic of Auto-SynPAR_Gi_, with activation of Gi read out by the TRUPATH BRET-based assay. Luc, NanoLuc luciferase. **(D)** TEV_cs_ and Thrombin_cs_-Auto-SynPAR_Gi_ constructs generate orthogonal, saturable, time- and dose-dependent activation of Gi in HEK293T cells as detected by TRUPATH. Data plotted as mean of triplicate, with error plotted as ±1 std. dev. This experiment was performed 2 times **(E**) Schematic of Auto-SynPAR_FL_, which can directly sense proteolysis in HEK293T cells by inducing enhanced fluorescence in a cpGFP inserted on the cytosolic side of the receptor. (**F)** TEV_cs_-Auto-SynPAR_FL_ responds specifically to TEVp treatment, but not Thrombin treatment. Data plotted as mean **Δ**F/F0 corrected for photobleaching for four regions of interest (quadrants) of the same field of view. Photobleaching correction was based on the addition of imaging buffer alone. Error is plotted as ±1 std. dev. This experiment was performed 3 times. **(G)** Selected images from data in (F) show un-corrected images at t=0, 5, and 15 minutes for TEV_cs_-Auto-SynPAR_FL_ treated with TEVp (top) or thrombin (bottom). From **Supplementary Movies 1-3**. Scale bars, 10 microns.

Next, we wanted to query the scope of outputs accessible to SynPARs beyond transgene expression. Our prior work on PAGER suggested that we would be able to engineer protease-dependent chemogenetic control of G-protein signaling by inserting a cleavage site between paired muscarinic toxins (MTs) and muscarinic acetylcholine receptor-based DREADDs.^14^ Indeed, we were able to achieve dose-dependent and orthogonal activation of Gi coupling in the TRUPATH assay, by adding either TEV_p_ or thrombin in the presence of the DREADD agonist deschloroclozapine (DCZ) (**Figure S5 A, B**).^15^ To extend the G-protein coupling output to the Auto-SynPAR scaffold we removed the SPARK fusion of Auto-SynPAR_TF_ and capitalized on the native Gi coupling of *k*OR. This allowed us to generate orthogonal TEV_p_ and Thrombin sensitive constructs that demonstrated both time- and concentration-dependent activation of Gi with the addition of protease (**Figure 2 C, D**).

The Auto-SynPAR scaffold was also amenable to creation of a real-time fluorescent biosensor for proteolysis. GPCRs have been converted into fluorescent biosensors for their ligands by inserting circularly permuted fluorescent proteins into conformationally sensitive locations within the GPCR of interest.^16^ Prior work on opioid sensors yielded *κ*Light 1.3, based on the *κ*OR scaffold, which rapidly increases fluorescence intensity in response to dynorphin binding.^17^ Because our Auto-SynPAR scaffold uses masked dynorphin, we posited that incorporation of *κ*Light 1.3 would yield a real-time protease sensor (**Figure 2E**). Indeed, we found that TEV_cs_-Auto-SynPAR_FL_ could specifically detect TEV protease activity on HEK293T cells via a 2 fold increase (**Δ**F/F0 ∼1) in green fluorescence (**Figure 2F, G**). In total these data support the use of SynPAR as a flexible, multimodal platform for studying protease activity *in situ*.

### SynPARs are amenable to library-based selections in mammalian cells

Our results from using published cleavage sequences to test the input modularity of SynPAR (Figure 2A, B) suggest that the performance of a cleavage sequence is a composite feature of protease activity for that sequence and the attributes of a SynPAR containing that cleavage site. This supports identifying and optimizing cleavage sequences *in situ* rather than grafting in sequences that are optimized *in vitro*. To this end we developed a method for mammalian-cell based selections of libraries of Chemo-SynPAR_TF_ constructs (**Figure 3A**). We generated cell lines for TEV_cs_ (ENLYFQ/GS) and Thrombin_cs_ (LVPR/GSGS) containing constructs and confirmed that they could respond appropriately when primed with the corresponding protease (**Figure 3B**). We then conducted model selections by mixing the TEV_cs_ and Thrombin_cs_ cell lines. Mixtures were primed and stimulated with TEV_p_ and sorted based on tBFP expression (**Figure 3C**). Genomic DNA (gDNA) was extracted and the region encoding the cleavage site was sequenced. (**Figure 3D, E, Figure S6A**). Based on these data we could expect to screen libraries with diversity ranging from 10^3^-10^4^. We also generated a pooled alanine scanning library for the TEV_cs_ and Thrombin_cs_ templates (**Figure S7A**). Cells were treated with either TEV_p_ or Thrombin prior to stimulation and analyzed as above. The data was consistent with established substrate specificities for both TEV_p_ and Thrombin. Unexpectedly, the P2’A mutant of the Thrombin_cs_ exhibited enhanced activity in the pooled screen. We characterized its activity in the luciferase reporter assay relative to the template Thrombin_cs_ using a thrombin dose response (**Figure S7B, C**). This experiment demonstrated that while the Thrombin EC_50_ was similar for the two constructs, the span of the P2’A curve was larger, suggesting that it may not be a better substrate, but rather a better SynPAR (**Figure S7D**). This highlights the importance of screening candidate cleavage sequences *in situ*.

**Figure 3.**
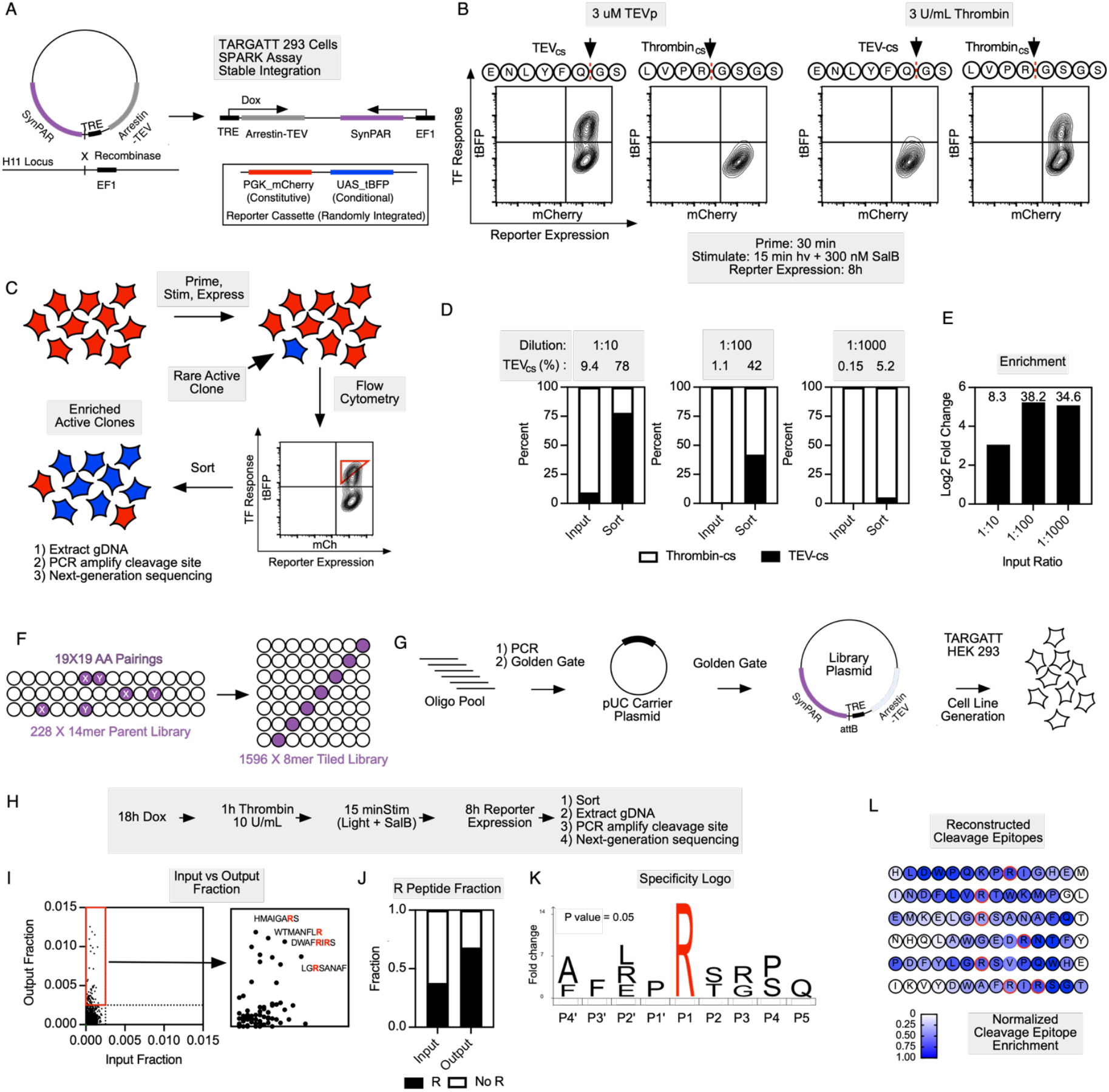
Chemo-SynPAR-TF enables library-based screens. **(A)** Stable single-copy integration of Chemo-SynPAR_TF_ and a Dox-inducible Arrestin-TEV using TARGATT expression system^20,21^. **(B)** TARGATT cell lines generated with TEV_cs_ and Thrombin_cs_ Chemo-SynPAR_TF_ respond specifically to TEVp and thrombin priming followed by coincident stimulation with 300 nM SalB and light for 15 min as detected by flow cytometry measuring reporter response. **(C)** Workflow for model selections of Chemo-SynPAR_TF_ expressed in TARGATT 293 cells to enrich rare active clones (blue) from a background of inactive cells (red) using FACS and next generation sequencing (NGS). **(D)** Results of model selections with 1:10, 1:100, and 1:1000 dilutions of TEV_cs_ TARGATT cells into Thrombin_cs_ TARGATT cells, demonstrating reliable enrichment of TEV_cs_ cells at all three dilutions. **(E)** Model selection enrichments. **(F)** The multiplexed substrate profiling library (MSP) is based on 228 14mer peptides comprising all XY, X_Y, and X_ _Y pairings of non-cysteine amino acids that has been tiled into 1596 8mer peptides. **(G)** The library is cloned from an oligo pool into a pUC carrier plasmid and then the TARGATT plasmid using sequential Golden Gate reactions prior to generation of the pooled library in the TARGATT 293 cell line. **(H)** The library was induced with doxycycline, primed with 10 U/mL of Thrombin for 1h, stimulated with ambient light and 300 nM SalB prior to reporter expression (8h), sorting, and processing for NGS. **(I)** Comparing the fractional composition of the input library and the output shows clear enrichment of peptides that contain arginine. **(J)** Arginine (R) containing peptides are enriched in the most abundant peptides in the output. **(K)** iceLogo comparing the arginine-adjacent residues of cleaved sequences (Output fraction > .0025) to all arginine-adjacent residues recapitulates known secondary determinants of Thrombin cleavage.^22^ **(L)** Reconstruction of overlapping peptides from the parent 14mer peptide enables the identification of cleavage epitopes by averaging the enrichment of all peptides that contain a given residue.

To expand our SynPAR libraries, we adapted peptide libraries designed for Multiplexed Substrate Profiling by Mass Spectrometry (MSP-MS), which are designed to achieve maximal protease substrate diversity with minimal library size.^18,19^ We designed a library of 228 different 14mer peptides and broke them down into 1596 tiled 8mers (7 per 14mer scaffold). This library was introduced into the Chemo-SynPAR_TF_ scaffold, which was integrated at one copy per HEK 293T cell using the TARGATT system (**Figure 3F, G, Figure S8**).^20,21^ We evaluated the library using Thrombin, a highly specific human protease. As such, we would not expect the ideal substrate for thrombin to be present in our library and must instead rely on analysis of the ensemble of cleaved sequences to reconstitute key determinants of cleavage. Comparison of the fractional representation of all library members in the input and output samples suggested specific enrichment (**Figure 3H, I, Figure S6C, Table S1**). We drew an arbitrary cutoff (0.0025) such that all the enriched sequences in the output would exceed the fractional representation of the most abundant sequence in the input. Thrombin is highly specific for P1 Arg, and when we examined the sequences that are most enriched, we observed that they all contain at least one Arg residue. Library-wide, peptides enriched in the output contain Arg with a higher frequency (68%) than in the input (37%) (**Figure 3J**). In addition, by comparing the representation of amino acids proximal to Arg sites that are cleaved (Output fraction >.0025) relative to all the Arg proximal amnio acids in the library, we enrich known secondary determinants of Thrombin activity, namely the P2 proline and small residues (Ser and Thr) at the P1’ position (**Figure 3K**).^22^ The structure of the library also allows the extraction of information about the determinants of cleavage among related sequences. Each cleaved sequence belongs to a family of seven overlapping peptides. By identifying parent 14mer peptides for which >3 members were highly enriched (Output fraction >.0025) and extracting the data for the remaining members of the family, we can reconstruct cleavage epitopes and better understand the determinants of SynPAR function (**Figure 3L**). The MSP-based library provides access to rich data and is a tractable platform for SynPAR reprogramming and optimization, as well as discovery of new protease substrate sequence determinants.

### Auto-SynPAR_Gi_ enables protease-dependent silencing of neurons in culture and *in vivo*

Chemogenetic control complicates the application of synthetic receptors in genetic or cell-based medicines. By not requiring an exogenous agonist, Auto-SynPAR overcomes this limitation and can be programmed to respond autonomously to protease activities that contribute to disease states. Tryptic protease activity plays an important role in inflammatory pain.^23^ Classically, proteases such as tryptase and kallikreins are secreted by immune cells that are recruited to sites of injury. These proteases activate PAR2 by cleaving its N-terminus to unmask a latent agonist.^24^ PAR2 couples predominantly through Gq/G12 in peripheral nociceptors, leading to hypersensitization and hyperalgesia. Endogenous and exogenous opiates can counteract this sensitization by activating Gi signaling, opening GIRK channels, hyperpolarizing nociceptors, and producing analgesia (**Figure 4A**).^25^ Since Auto-SynPAR_Gi_ is a protease-gated *κ*OR, we wondered whether it could create a synthetic, spatially constrained, and temporally resolved analgesic response at sites where protease activates endogenous PAR2. Such an approach would be more specific and homeostatic than using DREADDs or systemic opiates to silence nociceptors.

**Figure 4.**
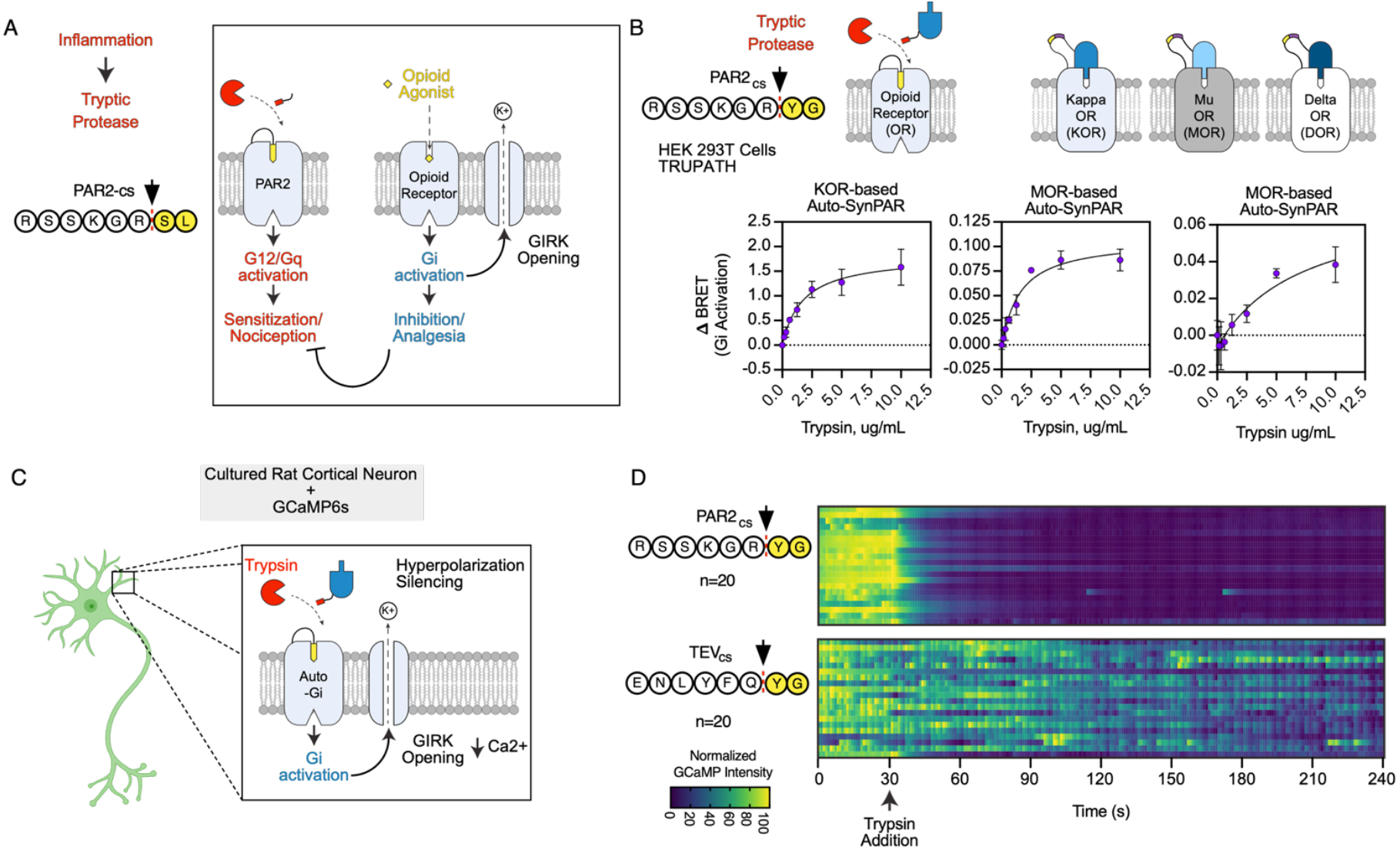
Auto-SynPAR_Gi_ can silence neurons in response to tryptic protease activity. **(A)** Endogenous protease-activated receptor 2 (PAR2) is activated by tryptic proteases that are secreted during inflammation. PAR2 couples to Gq and G12 signaling pathways in peripheral sensory neurons, leading to sensitization and enhanced nociception. Endogenous Opioid Receptors (ORs) inhibit these nociceptors by responding to agonists that couple to Gi signaling pathways. (**B**) *κ*, μ, and δ opioid receptors can be converted to Auto-SynPAR_Gi_ constructs that respond to Tryptic proteases. Receptors are fused to an antagonist via a linker containing the DynA_1-11_ sequence masked with the PAR2_cs_ (RSSKGR/DynA_1-11_). TRUPATH shows dose-dependent Gi activation in HEK 293T cells in response to trypsin. Data plotted as mean of triplicate, with error plotted as ±1 std. dev. This experiment was performed 2 times **(C)** Schematic of cultured rat cortical neuron expressing the calcium sensor GCaMP6 and a *k*OR-based PAR2_cs_-Auto-SynPAR-Gi construct to facilitate protease-specific silencing. GIRK, G-protein coupled inwardly rectifying potassium channel. **(D)** Plots of normalized GCaMP intensities over time, from rat cortical neurons expressing PAR2_cs_-Auto-SynPAR_Gi_ (top) or TEV_cs_-Auto-SynPAR_Gi_ (bottom). Treatment of these neurons with Trypsin protease (5 ug/mL at t=30s, arrow) at DIV 14 was sufficient to specifically silence spontaneous firing in neurons expressing the matched PAR2_cs_-Auto-SynPAR construct. This experiment was performed 3 times. See **Supplementary Movies 6, 7**.

To test this idea, we first introduced the PAR2_cs_ (RSSKGR) into the Auto-SynPAR_Gi_ scaffold and confirmed that we could generate trypsin-dependent Gi coupling in the TRUPATH assay. Since dynorphin can also activate μOR and δOR, we generated Auto-SynPAR_Gi_ constructs based on these GPCR scaffolds using a previously described inhibitory nanobody (NbE) in conjunction with a nested dynorphin_1-11_ agonist.^26^ These μOR and δOR derived constructs were also capable of generating protease-dependent Gi coupling, providing us access to a full complement of Auto-SynPARs based on the classical opioid receptors (**Figure 4B, Figure S9**). To establish protease-dependent silencing activity for *κ*OR-based Auto-SynPAR_Gi_, we transduced cultured rat cortical neurons with GCaMP6s and either TEV_cs_ or Thrombin_cs_-Auto-SynPAR_Gi_.^27^ Robust silencing was observed after 1 minute treatment with 1 uM TEV_p_ in neurons expressing TEV_cs_-Auto-SynPAR_Gi_. Conversely, no silencing was observed in those expressing the Thrombin_cs_ construct, supporting our ability to specifically modify neuronal activity in response to proteolysis (**Figure S10, Figure S11A Supplementary Movies 4, 5**). Next, we used the GCaMP assay described above to follow spontaneous firing in cultured neurons expressing either *κ*OR-based PAR2_cs_-Auto-SynPAR_Gi_ or TEV_cs_-Auto-SynPAR_Gi_ (**Figure 4C**). These constructs were generated with a P2A_mScarlet to enable assessment of expression by imaging (**Figure S11B, C**). We observed instantaneous silencing upon the addition of 5ug/mL Trypsin in cells expressing the PAR2_cs_ and as expected, saw no effect on spontaneous firing in cells expressing the TEV_cs_ (**Figure 4D**). This supports our ability to specifically silence neurons in response to a typically sensitizing perturbation.

To demonstrate the potential of PAR2_cs_-Auto-SynPAR_Gi_ to counteract a tryptic nociceptive stimulus *in vivo* we leveraged a *Scn10a*-Cre mouse line that enables targeted expression in the Dorsal Root Ganglion (DRG) and peripheral nociceptors when delivered by intrathecal injection of AAV-PHP.S.^28^ An intraplantar injection of trypsin was used to expose the nerve endings to a PAR2-activating stimulus that simultaneously activates an Auto-SynPAR_Gi_ (**Figure 5A**). Mechanical hyperalgesia was assessed by the von-Frey Filament (VFF) test (**Figure 5B**).^29^ The VFF model measures the withdrawal threshold for the trypsin-injected paw when stimulated with a filament of a given tensile strength. We injected mice with virus encoding either PAR2_cs_-Auto-SynPAR_Gi_ or TEV_cs_-Auto-SynPAR_Gi_, each with a P2A mScarlet, and waited 14 days for the gene to integrate and express. Expression in the DRG was assessed by detection of mScarlet and staining with anti-ALFA-647. Fluorescence microscopy of the stained DRG cross-sections suggested that both constructs were specifically expressed in the target population of neurons (**Figure 5C**). Prior to injection, animals from all groups had a withdrawal threshold of 1.2 g. We then measured the threshold for all three groups of mice at 1h, 2h and 3h after trypsin injection into the paw. We were able to observe an increase in withdrawal threshold with the PAR2_cs_ (n=4) relative to control mice expressing the TEV_cs_ (n=3) or no receptor (NR, n=2) (**Figure 5D, Figure S12A, B**). When we compare at the area under the curve (AUC) for withdrawal threshold at 1, 2, and 3 h timepoints we see a statistically significant difference between mice expressing the PAR2_cs_ and the TEV_cs_ (p=0.015) or NR (p=0.01) (**Figure 5E**). Therefore Auto-SynPAR_Gi_ can counteract pain signaling by converting a nociceptive proteolytic stimulus into analgesia. This technology represents a compelling prototype for a genetic medicine for pain that is activated locally and autonomously.

**Figure 5.**
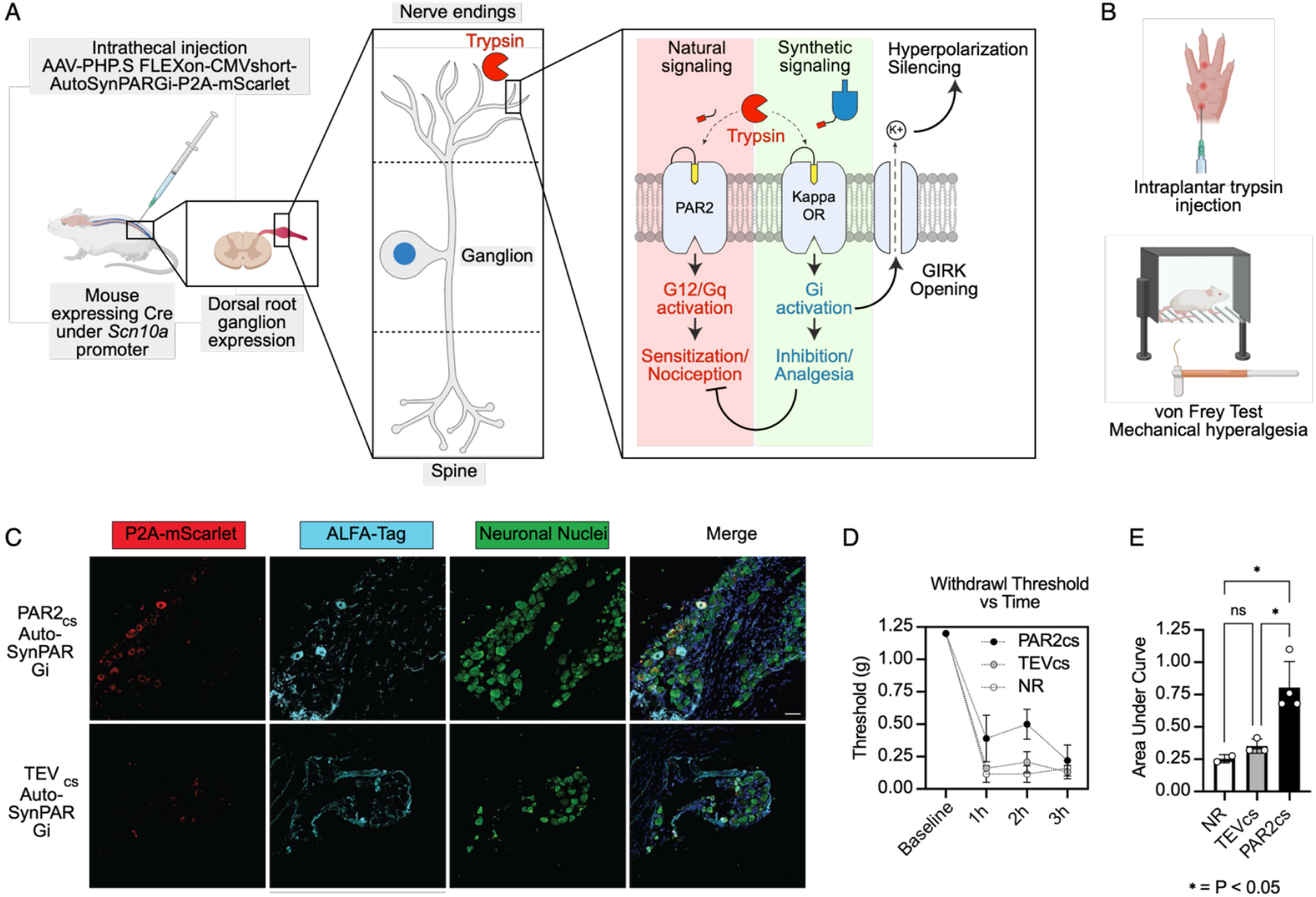
Auto-SynPARGi attenuates trypsin-induced mechanical hyperalgesia *in vivo*. **(A)** Mice expressing Cre under a *Scn10a* promoter were injected intrathecally (IT) with AAV-PHP.S encoding either PAR2_cs_- or TEV_cs_-Auto-SynPARGi with a P2A mScarlet to facilitate detection in the dorsal root ganglion (DRG) which innervates the extremities with nociceptors. Expression of Auto-SynPAR_Gi_ should counteract the pain induced by trypsin injection, which activates endogenous PAR2, by activating Gi and inducing neuron hyperpolarization. **(B)** To assess pain response, animals are subjected to an interplantar injection of trypsin and assessed using the von Frey Filament (VFF) test for paw withdrawal from filaments with standardized tensile strengths. **(C)** Imaging of Auto-SynPARGi expression in the DRG, 14 days after virus injection. Scale bar, 50 microns. **(D)** Results of VFF assay as in (B) measuring withdrawal threshold for the injected paw. Mice expressing PAR2_cs_-Auto-SynPARGi (n=4), TEV_cs_-Auto-SynPARGi (n=3), or neither (NR, n=2) were assessed 1, 2, and 3 h after trypsin injection. PAR2_cs_-expressing mice exhibited an increased withdrawal threshold relative to the control groups. **(E)** Quantification of area under the curve (AUC) across time points for data in (D).

## Discussion

The SynPAR platform provides an integrated toolset for developing genetically encoded recorders (SynPAR_TF_,), sensors (SynPAR_FL_), and actuators of cellular behavior (SynPAR_G_) in response to pericellular protease activity. SynPARs can function with chemogenetic control (Chemo-SynPAR) to enhance temporal specificity of the output function and provide the opportunity to integrate more proteolytic events prior to activation. Alternatively, SynPARs can be made to function autonomously (Auto-SynPAR) for simplified real-time sensing and use *in vivo*. We demonstrate input modularity using “off-the-shelf” cleavage sequences to reprogram input specificity and show that optimizing the cleavage sequence *in situ* using library-based selections may yield enhanced performance. Finally, we demonstrate that we can engineer an Auto-SynPAR_Gi_ to convert a proteolytic nociceptive stimulus into neuronal silencing and analgesia in culture and *in vivo*.

Extracellular protease activity has been implicated in a wide range of healthy and disease-associated signaling pathways. While some biological roles for protease activity are well established (coagulation, metastasis, nociception, etc.), emerging work suggests growing importance of extracellular protease signaling, such as in immunomodulation and regulation of host-pathogen interfaces.^30,31^ Importantly, these emerging niches for extracellular protease function are products of cell-cell interactions in complex tissues. This complexity necessitates technologies that can operate *in situ*, supporting the continued development of multimodal, genetically encoded, protease-sensitive tools like SynPARs.

In our work, we leveraged DREADD-based systems for Chemo-SynPAR (KORD for TF, hM4Di for Gi) and opioid receptors for Auto-SynPAR (*κ*OR for TF; KOR, μOR and δOR for Gi; *κ*OR-based *κ*-Light 1.3 for FL). These systems do not represent a full complement of input/output combinations, but rather a subset for which we identified immediate and robust use cases. Based on our work with PAGER, we anticipate that Chemo-SynPAR scaffolds that couple to Gs, Gq, G12, and FL should be readily accessible if desired. Generating Auto-SynPARs that couple to the remaining G protein families (beyond Gi) will likely require the use of other GPCRs that bind peptide agonists via an interaction with the N-terminus of the ligand.

Protease-responsive drugs, including conditionally active antibodies, cytokines, and T-cell engagers have gained popularity to limit on-target off-tissue effects of cytotoxic modalities in oncology.^32–34^ While these molecules have paved the way for protease-conditioned therapeutics, they have largely focused on conditional killing activities mediated by aberrant proteolysis in malignant tissue. Auto-SynPAR has the potential to be the prototype for a next generation of cellular and genetic medicines that respond to proteolysis with more sophisticated and biologically-driven mechanistic underpinnings. We demonstrate that we can use Auto-SynPAR to convert the nociceptive stimulus into analgesia in a model of inflammatory pain. This approach is appealing because it restricts the activity of the SynPAR only to cells that are exposed to the target stimulus, tuning the opioid response based on need and, in theory, limiting systemic off-target effects. More work is necessary to optimize the sensitivity, specificity, and delivery of these receptors prior to any pre-clinical development. However, SynPARs represent a first step in expanding the scope of accessible protease-responsive therapeutic modalities.

## Acknowledgements

A.Y.T. is grateful to the GPCR collaborative of the St. Jude Children’s Research Hospital, NIH (R01MN135934), Chan Zuckerberg Biohub – San Francisco, Wu Tsai Neurosciences Institute Neuro-omics Initiative, Phil and Penny Knight Initiative for Brain Resilience, and Stanford Bio-X for funding. We also thank the Fondo Italiano per la Scienza 2022-2023 (FIS-2023-03323 to F.D.L.)

## Author Contributions

M.R, N.A.K. and A.Y.T. conceived the project, designed and characterized SynPAR constructs. G.K and M.R designed and characterized μ and *δ*OR based Auto-SynPAR_Gi_ constructs. R.T and M.R. prepared neurons for *in vitro* GCaMP assays. R.L and N.W.B. designed and executed *in vivo* experiments. M.C. and F.D. designed and prepared AAV for *in vivo* experiments.

M.R and A.Y.T wrote the manuscript with input from all the authors.

## Materials and Methods

### Plasmid constructs and cloning

Constructs for transient expression in HEK293T cells were cloned into the pAAV vector with a CMV promoter. The pCDH lentiviral vector was used for stable cell line generation with Lentivirus unless otherwise noted. For all constructs, standard cloning procedures were used. PCR fragments were amplified using Q5 polymerase (NEB). Vectors were digested with NEB restriction enzymes and ligated to gel-purified PCR products using T4 ligation, Gibson, NEB HiFi, or Golden Gate assembly. Unless otherwise noted, ligated plasmids were introduced into competent XL1-Blue, NEB5-alpha, or NEB Stable bacteria via heat shock transformation.

### Cell lines

HEK293T cells were obtained from ATCC (tested negative for mycoplasma). TARGATT^™^ HEK293 cells were obtained from Applied StemCell (Milpitas, California). Cell lines were cultured as monolayers in complete growth medium: Dulbecco’s Modified Eagle Medium (DMEM, Corning) containing 4.5 g L^−1^ glucose and supplemented with 10% Fetal Bovine Serum (FBS, VWR), 1% (v/v) GlutaMAX (Gibco), and 1% (v/v) Penicillin-Streptomycin (Corning, 5,000 units mL ^−1^ of penicillin and 5,000 μg mL ^−1^ streptomycin). Cells were cultured at 37 °C under 5% CO_2_. For experimental assays, cells were grown in 6-well, 12-well, 24-well, or 96-well plates pretreated with 20 µg mL^−1^ human fibronectin (Millipore) for at least 10 min at 37 °C.

### HEK293T cell transient transfection

A 1 mg mL ^−1^ solution of PEI Max (Polysciences, 24765) was prepared for transient transfection as follows. Polyethylenimine (PEI, 500 mg) was added to 450 ml of Milli-Q H_2_0 in a 500 mL glass beaker while stirring with a stir bar. Concentrated HCL was added dropwise to the solution until the pH was less than 2.0. The PEI solution was stirred until PEI was dissolved (∼2–3 h). Concentrated NaOH was then added dropwise to the solution until the pH was 7.0. The volume of the solution was then adjusted to 500 mL, filter-sterilized through a 0.22-µm membrane, and frozen in aliquots at ^−^20 °C. Working stocks were kept at 4 °C for no more than 1 month.

For transient transfection, HEK293T cells were grown in 6-well, 12-well, or 24-well plates pretreated with 20 µg mL ^−1^ human fibronectin (Millipore) for at least 10 min at 37 °C. Cells were grown to a confluency of ∼70–90% prior to transfection. DNA transfection complexes were made by mixing DNA and 1 mg ml^−1^ PEI solution in serum-free DMEM at a 1 μg DNA: 5 μL PEI (1 mg mL ^−1^): 100 μL serum-free DMEM. Complexes were allowed to form for 20 min at room temperature. After 20 min, complexes were diluted in complete DMEM up to the growth volume per well size (2.5 ml for 6-well, 1 ml for 12-well, and 500 μL for 24-well). The entire well volume of the HEK293T cells was replaced with the diluted complexes and allowed to transfect cells at 37 °C for 5–24 h. Complete transfection protocols including amounts of DNA and length of transfection are described for each experiment below.

### Lentivirus generation

To generate lentivirus, HEK293T cells were cultured in T25 flasks and transfected at ∼70% confluency with 2.5 μg of the pCDH lentiviral transfer vector of interest and packaging plasmids psPAX2 (1.25 μg) and pMD2.g (1.25 μg) with 25 µl PEI (1 mg ml^−1^; Polysciences). Approximately 72 h post-transfection, the cell medium was collected and centrifuged for 5 min at 300 x *g* to remove cell debris. Medium containing lentivirus was used immediately for transduction or was aliquoted, flash-frozen in liquid nitrogen, and stored at –80 °C for later use. Frozen viral aliquots were thawed at 37 °C prior to infection. For pHR_UAS_tBFP_PGK_mCherry, virus was produced as above but using pCMVdR8.91instead of psPAX2. This virus was concentrated 25X with Lenti-X (Takara Bio) flash-frozen in liquid nitrogen and stored at –80 °C for later use.

### AAV1/2 generation

To generate AAV in supernatant, HEK293T cells were cultured in T25 plate and transfected at approximately 80% confluency in complete DMEM. Per each well, the AAV vector containing the gene of interest (900 ng) and AAV packaging/helper plasmids AAV1 (450 ng), AAV2 (450 ng), and DF6 (1800 ng) incubated with 25 μl PEI in 250 μl DMEM (serum free) were used for transfection in 5 mL complete DMEM. After 48 h the cell medium was collected and centrifuged for 5 min at 300 x *g* to remove cell debris. Medium containing AAV was used immediately for transduction or was aliquoted, flash-frozen in liquid nitrogen, and stored at –80 °C for later use. Frozen viral aliquots were thawed at 37 °C prior to infection. For hSyn_Auto-SynPARGi_P2A_mScarlet constructs, AAV1/2 was produced in a large scale (3× 15-cm plates) and purified using a HiTrap heparin column (GE Healthcare) as previously described.^35^

### Firefly luciferase reporter SynPAR_TF_ experiments

HEK293T cells were plated in human fibronectin-coated 6-well dishes at a density of 750,000 cells per well and allowed to grow overnight (∼18 h) at 37 °C until they reached ∼70–90% confluency. After ∼18 h, the cells were transfected with 350 ng of the indicated SynPAR_TF_ receptor plasmid, 100 ng of NanoLuc-β-arrestin2-TEVp plasmid, and 150 ng of UAS-Firefly Luciferase (FLuc) plasmid. Cells were transfected for 5 h at 37 °C. After transfection, cells from each well were lifted and resuspended in 6 ml of complete DMEM to make an ∼400,000 cells per ml single cell suspension, and 100 μL of cell suspension (∼40,000 cells) was plated per well in a human fibronectin-coated white, clear bottom 96-well plate in triplicate. Plates were wrapped in aluminum foil to protect them from light and incubated at 37 °C overnight (∼18 h). After ∼18 h, cells were cleaved with protease and stimulated. For half-life experiments cells were seeded at (∼20,000 cells) was plated per well in a human fibronectin-coated white, clear bottom 96-well plate in triplicate to prevent overgrowth in the longer experiment.

TEV_cs_-Chemo-SynPAR_TF_ and TEV_cs_-Auto-SynPAR_TF_ constructs were treated with TEV protease diluted in complete DMEM for the time and concentration indicated; For the modularity experiments in figure 2B Chemo-SynPAR_TF_ constructs were treated for 60 min in DMEM (serum free) at the indicated concentrations. For half-life experiments the interval between priming and stimulation was varied as indicated.

Stimulation was performed in a dark room with a red light source. For light-based Chemo-SynPARTF stimulations, constructs were stimulated with SalB and ambient room white light for 15 min. Priming solution was removed from the 96-well plate by flicking off and dabbing excess on a paper towel. To initiate stimulation, 100 μL stimulation solution was added to each well for a total of 15 min. For BRET-based stimulations 1X NanoLuc was added to the stimulation solution and white light was omitted. After 15 min, stimulation solution was removed, and 100 μL of complete DMEM was added back to each well. Plates were again wrapped in aluminum foil and placed in 37 °C incubator for 8 h. After 8 h post-stimulation, medium was removed, wells were washed once with 125 μL DPBS, and then 50 μl of 1x Bright-Glo (2x diluted 1:1 in DPBS; Promega) was added to each well and incubated for 1 min. After 1 min, firefly luciferase luminescence was measured using a Tecan Infinite M1000 Pro plate reader using the following parameters: 1,000 ms acquisition time, green-1 filter (520–570 nm), 25 °C linear shaking for 10 s.

### TRUPATH G-protein activation BRET assay

HEK293T cells were plated in human fibronectin-coated 6-well dishes at a density of 1,250,000 cells per well and allowed to adhere and grow for 2–4 h at 37 °C. After ∼2-4 h, the cells were transfected 1:1:1:1 with 250 ng of the indicated SynPAR_G_ receptor plasmid, 250 ng of the corresponding Gα_i_1-RLuc8, 250 ng of Gβ3 TRUPATH plasmid, and 250 ng Gγ9-GFP2 TRUPATH plasmid. Cells were incubated at 37°C and transfection was allowed to proceed for ∼20–24 h. After transfection, cells from each well were lifted and resuspended in 6 ml of complete DMEM to make an ∼200,000 cells per ml single cell suspension, and 100 μl of cell suspension (∼20,000 cells) was plated per well in a human fibronectin-coated white, clear bottom 96-well plate in triplicate. Plates were incubated at 37 °C for ∼20–24 h.

For Chemo-SynPAR_Gi_ cells were treated with the indicated concentration of protease for 30 min in DMEM (serum free) at 37 °C followed by stimulation with 1 nM DCZ and 10 μM CTZ400a (substrate for TRUPATH assay) diluted in HBSS + 20 mM HEPES for 5 min before reading out BRET. For Auto-SynPAR_Gi_ cells were treated with the indicated concentration of protease diluted in and 10 μM CTZ400a diluted in HBSS + 20 mM HEPES, read immediately and at the intervals indicated. BRET was read out using a Tecan Infinite M1000 Pro plate reader using the following parameters: filter 1 magenta (370 to 450 nm), 500 ms integration time; filter 2 green (510 to 540 nm), 500 ms integration time; 25 °C. Data are presented as Delta BRET.

### HEK293T stable cell line generation

HEK293T cells were plated on six-well human fibronectin-coated plates. When cells reached ∼70–90% confluency, cells were transduced with lentivirus for 1–3 days. After transduction, cells were expanded and frozen for later use without further selection.

### Preparation of SynPAR_TF_ MSP (multiplex substrate profiling) libraries

For the TEV/Thrombin alanine scanning library individual single stranded oligos were ordered from Elim Biopharmaceuticals (Hayward, California) flanked by PaqCI Golden Gate sites. The oligos were pooled and converted into double stranded DNA and amplified by PCR. This fragment was purified and used as the donor to be inserted into the TARGATT^™^ plasmid containing a SynPAR-TF receptor with ccdB and chloramphenicol kill cassette flanked by PaqCI Golden Gate sites in the position for cleavage site insertion. For the MSP library, a single stranded oligo pool containing the library flanked by PaqCI Golden Gate sites was ordered from Twist Bioscience (South San Francisco, California). The oligo pool was converted into double stranded DNA and amplified by PCR using primers containing an Esp3I Golden Gate site, purified, and subsequently used as the donor to be inserted into a pUC19 plasmid containing Esp3I Golden Gate sites flanking a ccdB and chloramphenicol kill cassette for counter selection. The product of the Esp3I Golden Gate reaction was purified and electroporated into NEB10-beta electrocompetent cells. Transformation efficiency was confirmed by serial plating to be at least 1000x library size. This created a pUC19 MSP cleavage site library flanked by PaqCI Golden Gate sites that could be stored and amplified in bacteria without compromising library integrity. This pUC19 MSP cleavage site library was then used as the donor to be inserted into the TARGATT^™^ plasmid containing SynPAR_TF_ receptor with ccdB and chloramphenicol kill cassette flanked by PaqCI Golden Gate sites in the position for cleavage site insertion.

The product of the PaqCI Golden Gate reaction was purified and electroporated into NEB10-beta electrocompetent cells. Transformation efficiency was confirmed by serial plating to be at least 1000x library size. This created a Chemo-SynPAR_TF_ TARGATT^™^ plasmid library that could be stored and amplified in bacteria without compromising library integrity. These Chemo-SynPAR_TF_ TARGATT^™^ libraries were then used as the donor for integration into the TARGATT^™^ HEK293 landing pad.

### TEV, Thrombin, and MSP SynPAR_TF_ stable cell line generation using TARGATT HEK293

TARGATT HEK293 cells are commercially available cells that contain a landing pad in the H11 locus of the genome that can be used to specifically integrate a transgene of interest at this site. The TEV_cs_ and Thrombin_cs_ Chemo-SynPAR_TF_ stable cell lines were generated using TARGATT HEK293 cells. The UAS_tBFP_PGK_mCherry reporter was randomly integrated into the genome of these cells, while the integrase landing pad was used to integrate the SynPAR receptor and the doxycycline inducible arrestin-TEVp components. Briefly, to integrate the reporter component of SynPAR_TF_, TARGATT HEK293 cells were plated on twelve-well human fibronectin-coated plates. When cells reached ∼70–90% confluency, cells were transduced with reporter lentivirus for 2–3 days. These cells were then lifted, scaled up, and frozen back for future use. Using these reporter-expressing TARGATT HEK293 cells, the receptor and doxycycline inducible arrestin-TEVp components of SynPAR-TF were integrated into the landing pad according to the manufacturer’s instructions. The cassette to be integrated also contained a puromycin marker for selection of successful integration. Briefly, the cells were plated on six-well human fibronectin-coated plates. When cells reached ∼70–90% confluency, cells were transfected with 500ng of integrase plasmid and 500ng of donor plasmid using PEI. Cells were transfected for 48 hours before they were then lifted and replated in a T75 flask. For the MSP library, 18 individual 6 well transfections were conducted in parallel to maximize library homogeneity. Three days later (5 days total post transfection), stably integrated and expressing cells were selected for in complete DMEM containing 0.5 μg ml^−1^ puromycin for 5 days. Cells were split, expanded and pooled ass appropriate when they reached ∼80–90% confluency. After the 5 day selection, puromycin was removed and cells were used for experiments or frozen for future use. Of note, decreased activation of the UAS-reporter was observed overtime, presumably due to promoter silencing. To overcome this, these cells were transduced with reporter lentivirus for a second time and subsequent experiments were conducted within 2 weeks of transduction.

### Cell sorting of SynPAR-TF library

SynPAR-TF stable cells contain a constitutively expressed mCherry fluorescent marker, a constitutively expressed Chemo-SynPAR_TF_ receptor, a doxycycline-inducible arrestin-TEVp, and a UAS-tBFP reporter that expressed only if Chemo-SynPAR_TF_ is activated. To activate them, SynPAR-TF stable cells were plated such that they would be ∼80% confluent after 24 hours. At the time of plating, 10 μg/mL doxycycline was added to the growth media and the plate/flask was wrapped in aluminum foil to protect from light. Cells were allowed to grow and arrestin-TEVp be induced for 24 hours. After 24 hours of induction, and in the dark under red light, the media was removed from the cells, fresh serum-free media containing protease was added to the cells, and the cells were allowed to incubate in the dark at 37 °C for 1 hour. After 1 hour, in the dark and under red light, the media was removed and fresh complete growth media containing 300 nM Sal B was added to the cells. The cells were then incubated under white light and at room temperature for 15 minutes. After the 15 min stimulation, in the dark and under red light again, the media was removed and replaced with complete growth media. The plate/flask was again wrapped in aluminum foil and incubated at 37 °C for 8 hours before processing and cell sorting.

SynPAR-TF stable cells were sorted on a Sony SH800 Cell Sorter using a 100-μm microfluidic sorting chip. Briefly, ∼8 hours post stimulation, SynPAR-TF cells were lifted using an enzyme-free dissociation buffer (Thermo 13151014), washed once with FACS buffer (PBS + 3% FBS), and kept on ice until sorting. A control sample that was stimulated with SalB but not treated with protease was used to set the sorting gate. Activated SynPAR-TF cells (mCherry+ and BFP+ double positive cells) were cells sorted into complete growth medium and kept on ice for downstream processing and sequencing. Sorted cells were pelleted and gDNA extracted using DNeasy Blood and Tissue Kit (Qiagen) per the manufacturer’s instructions.

gDNA was concentrated to 10 uL using a DNA Clean and Concentrator Kit (Zymo Research). 20-100% of the extracted and concentrated gDNA was used as template to generate a PCR amplicon spanning the cleavage site for NGS. To enhance specificity PCR reactions were run in 1 mM Betane using a Touchdown PCR method with an annealing temperature starting at 64 °C and dropping 0.4 °C per cycle for 30 cycles. Amplicons were cleaned up using a DNA Clean and Concentrator Kit, concentration was measured using a Qubit, and samples were sent for either miSeq (TEV/Thrombin alanine scanning library) or Nanopore (Model & MSP selections) analysis.

### Confocal Microscope Hardware and Software

Confocal imaging was performed on a Zeiss AxioObserver inverted confocal microscope with 10× and 20× air objectives, and 40× and 63× oil-immersion objectives, outfitted with a Yokogawa spinning disk confocal head, a Quad-band notch dichroic mirror (405/488/568/647), and 405 (diode), 491 (DPSS), 561 (DPSS) and 640 nm (diode) lasers (all 50 mW). The following combinations of laser excitation and emission filters were used for various fluorophores: GFP (491 laser excitation; 528/38 emission), mCherry/Alexa Fluor 568 (561 laser excitation; 617/73 emission), Alexa Fluor 647 (647 excitation; 680/30 emission), and differential interference contrast. Acquisition times ranged from 100 ms. All images were collected using SlideBook (Intelligent Imaging Innovations) and processed using FIJI/ImageJ.

### Imaging SynPAR_TF_ localization

HEK 293T cells were transfected as if for a SynPAR_TF_ firefly luciferase experiment (above). After transfection cells were plated on glass coverslips coated with human fibronectin, wrapped in foil, and grown overnight. Cells were primed and stimulated as indicated. Immediately after stimulation, cells were fixed in 4% paraformaldehyde for 10 min at room temperature, followed by membrane permeabilization in ice cold methanol for 10 min. The cells were then incubated in PBS supplemented with 1% BSA for 30 min for blocking, followed by 1:1,000 anti-ALFA–AlexaFluor647 and 1 µg ml^−1^ DAPI in blocking buffer for 1 h to stain for SynPAR_TF_ localization with a nuclear marker. The cover slips were then mounted on slides and analyzed by confocal microscopy.

### Fluorescence imaging of SynPAR_**FL**_

35 mm glass bottom dishes (CellVis) were plates pretreated with 20 µg mL ^−1^ human fibronectin (Millipore) and seeded with cells stably expressing TEV_cs_-Auto-SynPAR_FL_ ∼200,000 cells/mL and allowed to grow for 18-24h.Cells were washed 2 × 2 mL with HBSS (+Magnesium Chloride, +Calcium Chloride), and then incubated in 100 ul of the same. Images of GFP intensity were captured at every 0.25 min. After 1.5 min, 100 uL of 2X protease solution was added to the dish. Images were acquired for 240 s total. To account for photobleaching a sample with without protease was included.

Images were analyzed using FIJI/ImageJ software. The image was divided into 4 quadrants. Background was subtracted based on an ROI without cells. **Δ**F/F0 was calculated for each frame at in all 4 quadrants to yield a mean **Δ**F/F0 and error across the field of view.

### GCaMP6s neuronal activity assay

All procedures were approved and carried out in compliance with the Stanford University Administrative Panel on Laboratory Animal Care, and all experiments were performed in accordance with relevant guidelines and regulations. Before dissection, 35 mm glass bottom dishes (CellVis) were coated with 0.001% (w/v) poly-L-ornithine (Sigma-Aldrich) in DPBS (Gibco) at room temperature overnight, washed three times with DPBS, and subsequently coated with 5 μg mL^−1^ of mouse laminin (Gibco) in DPBS at 37 °C overnight. Cortical neurons were extracted from embryonic day 18 Sprague Dawley rat embryos (Charles River Laboratories, strain 400) by dissociation in Hank’s balanced salt solution with calcium and magnesium (Gibco). Cortical tissue was digested in papain according to the manufacturer’s protocol (Worthington), then 5 × 10^5^ cells were plated onto each dish in neuronal culture medium at 37 °C under 5% CO_2_. The neuronal culture medium is neurobasal (Gibco) supplemented with 2% (v/v) B27 supplement (Life Technologies), 0.5% (v/v) fetal bovine serum, 1% (v/v) GlutaMAX, 1% (v/v) penicillin-streptomycin, and 1% (v/v) sodium pyruvate (Gibco, 100 mM).

On DIV 6, half of the medium was removed from each dish and replaced with neuronal culture medium. On division 6 after the medium change, each well was infected with 25 μl of AAV1/2 (12.5 μl of GCaMP6s AAV and 12.5 μl of Auto-SynPAR_Gi_ AAV). Neurons were wrapped in aluminum foil and allowed to express in the incubator.

To check expression on DIV 13, cells were washed 2 × 2 mL with HBSS (+Magnesium Chloride, +Calcium Chloride), and then incubated in 100 ul 1:1,000 anti-ALFA–AlexaFluor647 for 3 min. Cells were then located under the microscope and time-lapse images imaged.

For silencing experiments cells were washed as above and incubated with 100 uL HBSS (+Magnesium Chloride, +Calcium Chloride). Images of GCaMP intensity were captured at 1 Hz. After 30 s, 100 uL of 2X protease solution was added to the dish. Images were acquired for 240 s total.

Images were analyzed using FIJI/ImageJ software. ROIs for individual neurons were manually added to images and the normalized GCaMP intensity for each neuron was plotted vs time.

### AAV-PHP.S purification for use *in vivo*

AAVPro-HEK293T cells were cultured in DMEM supplemented with heat-inactivated FBS (10 %), Pen/Strep (1%), and L-glutamine (2%) and sodium pyruvate (1%). Recombinant AAV particles (rAAVs) were produced by using triple transfection strategy as described previously.^36^ Three days before transfection, cells were seeded in a CellBIND Polystyrene CellSTACK 2 Chamber (#3310, Corning, Corning, NY, USA) for 48 h to reach 80% confluence. Cells were washed with PBS, detached with trypsin-EDTA 0.05% (EuroClone, Milan, Italy). Finally, 250 million cells were seeded in a and CellBIND Polystyrene CellSTACK 5 Chamber (#3311; Corning) for 24 h, until 80% of confluence. Each chamber was transfected with 2.5 mg of DNA containing the three plasmids (packaging, helper and plasmids expressing gene of interest) in a 1:1:1 molar ratio. As packaging plasmid was used AAV2 rep-AAV-PHP.S (pUCmini-iCAP-PHP.S #103006, Addgene), and as helper plasmid was used pAdDeltaF6 (#112867, Addgene). Total DNA was diluted in OptiMEM (Thermo Fisher Scientific) and used for cell transfection. Cells were left at 37°C for 72 h, before rAAVs collection. Sample was centrifuged at 3.200 × g for 15 min at 4 °C and supernatant, containing the AAV particles was collected and incubated with Benzonase (50 UI/mL; #E1014, Merck Millipore) at 37 °C for 45 min to digest residual plasmids and residual genomic DNA/cellular RNA. rAAVs were purified using a gradient of iodixanol as previously reported. Briefly, a gradient of iodixanol (OptiPrep; STEMCELL Technologies, Vancouver, Canada) was prepared (15%, 40% and 60%). Each solution was added into a 39-mL Quick-Seal tube (Beckman Coulter, Brea, CA, USA) using a syringe equipped with an 18 G needle. Finally, tubes were filled with the sample, sealed and, centrifuged in a Type 70 Ti rotor (Beckman Coulter) at 350.000 x g at 10°C for 90 min. Viral particles contained in the 40% iodixanol layer were fractioned in 1.5 mL microcentrifuge tubes and concentrated using Amicon Ultra-15 centrifugal filter units (molecular weight cut-off, 100 kDa; Merck Millipore). The viral titer was quantified using RT-qPCR using RT-qPCR using AAVpro® Titration Kit (#6233 Takara Bio).

### Animals

Male *Scn10a Na*_*v*_*1*.*8-Cre* mice (10 weeks old; JAX® #036564) were used for all experiments. This transgenic line carries Cre recombinase under the control of the *Scn10a* (sodium channel, voltage-gated, type X, alpha or Na_v_1.8) promoter. Animals were group housed in individually ventilated cages under standard laboratory conditions, including a 12-hour light/dark cycle (lights on at 7:00 AM), an ambient temperature of 22 ± 2 °C, and 40–60% humidity. Mice had *ad libitum* access to standard chow and water. All procedures involving animals were conducted in compliance with institutional guidelines and were approved by NYU Langone Institutional Animal Care and Use Committee (IACUC).

### Experimental Design

To achieve cell-type–specific expression of exogenous genes in nociceptors, we employed *Scn10a Na*_*v*_*1*.*8-Cre* mice. In these animals, Cre recombinase is selectively expressed under the control of the *Scn10a* promoter, which is active primarily in sensory neurons, including small-diameter dorsal root ganglion (DRG) neurons. This genetic specificity allows for targeted manipulation of nociceptors while sparing non-nociceptive neurons.

For Cre-dependent transgene delivery, we used an adeno-associated viral vector (*AAV2/php*.*s-CMVshort-PAR2-T2A-mScarlet*) designed to express both synPAR2 or TevPAR2 and the fluorescent reporter mScarlet. The transgene cassette is arranged such that expression is dependent on Cre-mediated recombination, ensuring that only *Nav1*.*8*-expressing neurons activate transgene expression. This strategy enables both functional manipulation of nociceptors and visual confirmation of viral transduction via mScarlet fluorescence.

All animals were initially acclimated to the behavioral testing environment, followed by a baseline assessment of mechanical sensitivity using the paw von Frey test. Following baseline testing, mice were randomly assigned to one of three experimental groups and received an intrathecal injection (i.t., 5 µL) of either:

- AAV2/php.s-CMVshort-PAR2-T2A-mScarlet (2 × 1012 vg/mL),
- AAV2/php.s-CMVshort-TEV.cs-T2A-mScarlet (>2 × 1012 vg/mL)
- Vehicle

Intrathecal injections were performed using a 10 µL Hamilton syringe under isoflurane anesthesia. Following viral delivery, mice were returned to their home cages and monitored daily. Fourteen days post-injection are required for adequate viral transgene expression and its potential downstream physiological effects. Seven days from injection, mice were re-acclimated to the behavioral setup and re-tested to obtain post-transduction von Frey baselines. On the experimental day, mice received an intraplantar (i.pl) injection of trypsin (140 nM in 10 µL of sterile saline) into the left hindpaw under brief isoflurane anesthesia. Mechanical sensitivity was then assessed using the modified up-and-down von Frey method hourly for 4 hours post-injection.

### Mechanical Sensitivity: paw von Frey Test

Mechanical sensitivity was assessed using the modified up-and-down von Frey method applied to the plantar surface of the hindpaw. Mice were placed individually in transparent Plexiglas chambers (10 × 10 × 15 cm) positioned on an elevated wire mesh platform, allowing access to the plantar surface of the hindpaws. Animals were acclimated to the testing environment for 2 hours for 3 consecutive days before baseline (on day 4) and experimental testing (on day 5).

A series of calibrated von Frey monofilaments (range: 0.007 to 2.0 g; Stoelting, USA) were applied perpendicularly to the center of the left hindpaw plantar surface with sufficient force to bend the filament for approximately 2-3 seconds. The modified up-and-down method was employed as described by Chaplan et al. (1994), with testing initiated using the g/force filament. A positive response was defined as a sharp paw withdrawal, shaking, or licking upon stimulus application. If the response was negative, a new, higher gram force filament was applied, continuing to increase filament’s gram force until a response was achieved. Then, an up (higher gram force filament) and down (lower gram force filament) was applied to the hindpaw to confirm reading accuracy. The paw withdrawal threshold was then calculated. To minimize experimenter bias, all behavioral assessments were performed by individuals blinded to experimental group assignments.

### Immunofluorescence and Confocal Imaging of animal tissue

Mice were deeply anesthetized and transcardially perfused with phosphate-buffered saline (PBS) followed by 4% paraformaldehyde (PFA) in PBS. Lumbar dorsal root ganglia (DRG; L1–L5) were dissected, post-fixed in 4% PFA for 2 hours at 4 °C, then cryoprotected in 30% sucrose overnight. Tissues were embedded in Tissue Freezing Media (General Data) compound and cryosectioned at 12 μm thickness onto Superfrost Plus slides. Sections were rinsed in PBS and blocked for 1 hour at room temperature (RT) with 10% normal donkey serum (NDS) in PBS containing 0.3% Triton X-100. Primary antibody incubation was carried out overnight at 4 °C in PBS with 3% NDS and 0.3% Triton X-100, using the following antibodies: Guinea pig anti-NeuN (1:500; Millipore, ABN90) to label neurons, Rabbit anti-α-tag (1:300; Cell Signaling Technology, #549637) to detect α-tagged receptor expression.

The following day, sections were washed in PBS (3 × 5 minutes) and incubated with species-appropriate Alexa Fluor-conjugated secondary antibodies (Goat anti Guine Pig Alexa 488 and Donkey anti Mouse Alexa 647; 1:1000) for 45 minutes at RT. After three final washes in PBS, sections were counterstained with DAPI (nuclear staining) and mounted using ProLong® Gold Antifade Mountant (Thermo Fisher).

Images were acquired using a Leica SP8 confocal microscope with sequential scanning settings to minimize spectral overlap. mScarlet fluorescence was visualized directly without the need for antibody amplification. Identical acquisition settings were used across experimental groups for quantitative comparisons.

### Quantification and Data Analysis

All graphs were created using GraphPad Prism 10. Error bars represent SD unless otherwise noted. For scatter plots, variable slope (four parameter) non-linear regression lines were used. For comparison between two groups, *P* values were determined using two-tailed Student’s *t*-tests. For multiple comparisons, *P* values were determined using two-way ANOVA with Tukey’s multiple comparisons test to adjust for multiple comparisons. \**P* < 0.05; \*\**P* < 0.01; \*\*\**P* < 0.001; \*\*\*\**P* < 0.0001; NS, not significant.

## References

1. Overall, C. M. & Blobel, C. P. In search of partners: linking extracellular proteases to substrates. Nat. Rev. Mol. Cell Biol. 8, 245–257 (2007).

2. Frei, M. S., Mehta, S. & Zhang, J. Next-Generation Genetically Encoded Fluorescent Biosensors Illuminate Cell Signaling and Metabolism. Annu. Rev. Biophys. 53, 275–297 (2024).

3. Sanman, L. E. & Bogyo, M. Activity-based profiling of proteases. Annu. Rev. Biochem. 83, 249–273 (2014).

4. Teng, F. et al. Programmable synthetic receptors: the next-generation of cell and gene therapies. Signal Transduct. Target. Ther. 9, (2024).

5. Peach, C. J., Edgington-Mitchell, L. E., Bunnett, N. W. & Schmidt, B. L. Protease-Activated Receptors in Health and Disease. Physiol. Rev. 103, 717–785 (2023).

6. Kalogriopoulos, N. A. et al. Synthetic GPCRs for programmable sensing and control of cell behaviour. Nature 637, 230–239 (2025).

7. Vardy, E. et al. A New DREADD Facilitates the Multiplexed Chemogenetic Interrogation of Behavior. Neuron 86, 936–946 (2015).

8. Kim, M. W. et al. Time-gated detection of protein-protein interactions with transcriptional readout. Elife 6, 1–24 (2017).

9. Roth, B. L. DREADDs for Neuroscientists. Neuron 89, 683–694 (2016).

10. Wang, Y. et al. Structures of the entire human opioid receptor family. Cell 186, 413–427.e17 (2023).

11. Lopez, M. I. S. & Ting, A. Y. Directed evolution improves the catalytic efficiency of TEV protease. bioRxiv 811570 (2019). doi:10.1101/811570

12. Vasiljeva, O., Menendez, E., Nguyen, M., Craik, C. S. & Michael Kavanaugh, W. Monitoring protease activity in biological tissues using antibody prodrugs as sensing probes. Sci. Rep. 10, 1–10 (2020).

13. Kim, C. K., Cho, K. F., Kim, M. W. & Ting, A. Y. Luciferase-LOV BRET enables versatile and specific transcriptional readout of cellular protein-protein interactions. Elife 8, 1–21 (2019).

14. Armbruster, B. N., Li, X., Pausch, M. H., Herlitze, S. & Roth, B. L. Evolving the lock to fit the key to create a family of G protein-coupled receptors potently activated by an inert ligand. Proc. Natl. Acad. Sci. U. S. A. 104, 5163–5168 (2007).

15. DiBerto, J. F., Olsen, R. H. J. & Roth, B. L. TRUPATH: An Open-Source Biosensor Platform for Interrogating the GPCR Transducerome. Methods Mol. Biol. 2525, 185–195 (2022).

16. Jing, M. et al. An optimized acetylcholine sensor for monitoring in vivo cholinergic activity. Nat. Methods 17, 1139–1146 (2020).

17. Dong, C. et al. Unlocking opioid neuropeptide dynamics with genetically encoded biosensors. Nat. Neurosci. 27, 1844–1857 (2024).

18. Donoghue, A. J. O. et al. Global identification of peptidase specificity by multiplex substrate profiling. Nat. Methods 9, 1095–1100 (2012).

19. Multiplex substrate profiling by mass spectrometry for proteases. in 375–411 (2023). doi:10.1016/bs.mie.2022.09.009

20. Chi, X., Zheng, Q., Jiang, R., Chen-Tsai, R. Y. & Kong, L.-J. A system for site-specific integration of transgenes in mammalian cells. PLoS One 14, e0219842 (2019).

21. Zhu, F. et al. DICE, an efficient system for iterative genomic editing in human pluripotent stem cells. Nucleic Acids Res. 42, e34–e34 (2014).

22. Colaert, N., Helsens, K., Martens, L., Vandekerckhove, J. & Gevaert, K. Improved visualization of protein consensus sequences by iceLogo. Nat. Methods 6, 786–787 (2009).

23. Steinhoff, M. et al. Agonists of proteinase-activated receptor 2 induce inflammation by a neurogenic mechanism. Nat. Med. 6, 151–158 (2000).

24. Stefansson, K. et al. Activation of Proteinase-Activated Receptor-2 by Human Kallikrein-Related Peptidases. J. Invest. Dermatol. 128, 18–25 (2008).

25. Nockemann, D. et al. The K + channel GIRK2 is both necessary and sufficient for peripheral opioid-mediated analgesia. EMBO Mol. Med. 5, 1263–1277 (2013).

26. Yu, J. et al. Structural basis of μ-opioid receptor targeting by a nanobody antagonist. Nat. Commun. 15, (2024).

27. Chen, T.-W. et al. Ultrasensitive fluorescent proteins for imaging neuronal activity. Nature 499, 295–300 (2013).

28. Jimenez-Vargas, N. N. et al. Protease-activated receptor-2 in endosomes signals persistent pain of irritable bowel syndrome. Proc. Natl. Acad. Sci. 115, (2018).

29. Chaplan, S. R., Bach, F. W., Pogrel, J. W., Chung, J. M. & Yaksh, T. L. Quantitative assessment of tactile allodynia in the rat paw. J. Neurosci. Methods 53, 55–63 (1994).

30. Donado, C. A. et al. Granzyme K activates the entire complement cascade. Nature 641, 211–221 (2025).

31. Lakemeyer, M. et al. A <em>Bacteroides fragilis</em> protease activates host PAR2 to induce intestinal pain and inflammation. Cell Host Microbe 33, 1686–1702.e11 (2025).

32. Cao, W. et al. A next-generation anti-CTLA-4 probody mitigates toxicity and enhances anti-tumor immunity in mice. Nat. Commun. 16, 9029 (2025).

33. Mansurov, A. et al. Masking the immunotoxicity of interleukin-12 by fusing it with a domain of its receptor via a tumour-protease-cleavable linker. Nat. Biomed. Eng. 6, 819–829 (2022).

34. Cattaruzza, F. et al. Precision-activated T-cell engagers targeting HER2 or EGFR and CD3 mitigate on-target, off-tumor toxicity for immunotherapy in solid tumors. Nat. Cancer 4, 485–501 (2023).

35. Kim, C. K. et al. A Molecular Calcium Integrator Reveals a Striatal Cell Type Driving Aversion. Cell 183, 2003–2019.e16 (2020).

36. Titiz, M. et al. Schwann cell C5aR1 co-opts inflammasome NLRP1 to sustain pain in a mouse model of endometriosis. Nat. Commun. 15, 10142 (2024).

