## Supplementary Information for "Programming cell behavior with synthetic protease-activated receptors"

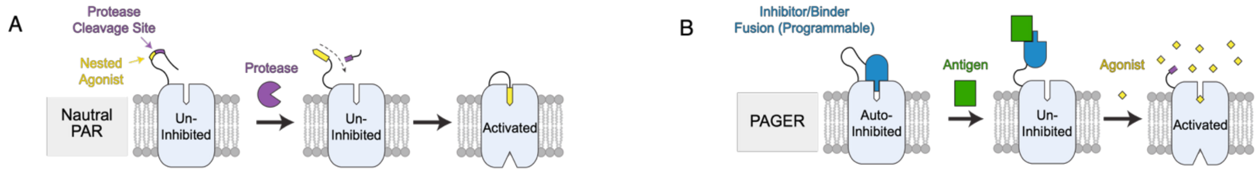


**<u>Supplementary Figure 1</u> Schematics for natural PARs and PAGERs. (A)** Natural PARs exist in an un-inhibited state. Protease cleavage unmasks a latent agonist that binds and activates the receptor in *cis*. **(B)** PAGERs are engineered GPCRs that are inhibited in *cis* by an inhibitor/binder fusion. Antigen binding sterically interferes with inhibitor to relieve inhibition, allowing receptor activation by an exogenous agonist.

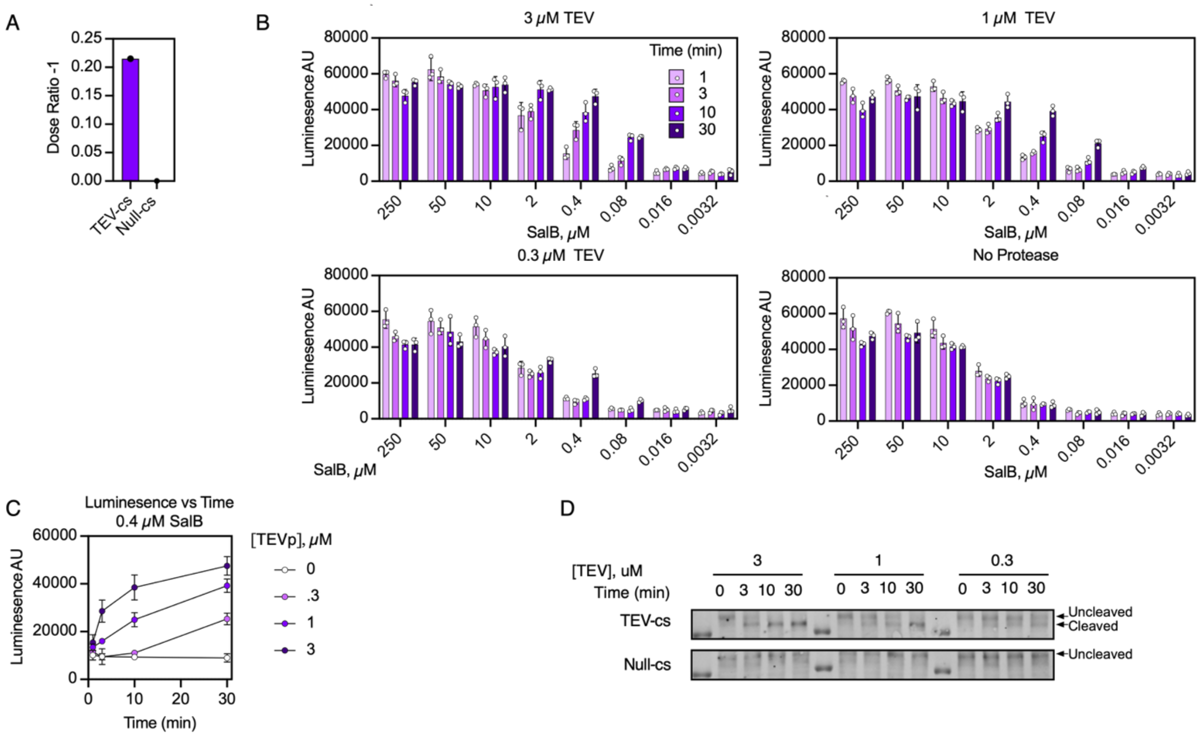


**Supplementary Figure 2 SynPARs integrate pericellular protease activity. (A)** Dose ratio -1 comparison for $TEV_{cs}$ and $Null_{cs}$ $SynPAR_{TF}$ constructs in Figure 1C. **(B)** Raw luminescence data for Figure 1D demonstrating the effect of agonist (SalB) concentration, time, and protease concentration on reporter signal. Mean intensities plotted as bars, individual values (n=3) plotted as circles, error bars are SD. **(C)** Plot of luminescence vs time for varying protease concentrations and times. Mean intensities plotted as circles; error bars are SD. **(D)** Western blot monitoring cleavage of $SynPAR_{TF}$ constructs at different protease and incubation times. All experiments are representative of at least biological duplicates.

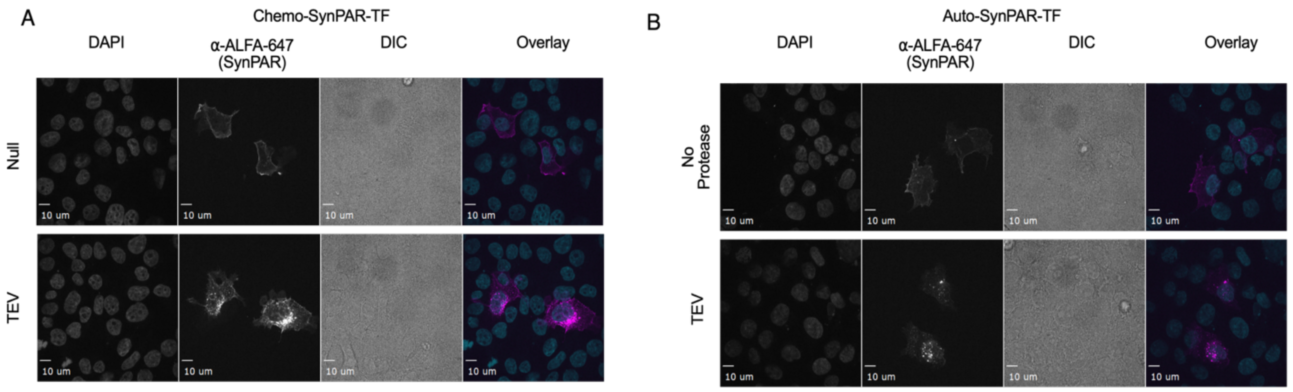


**<u>Supplementary Figure 3</u> Confocal microscopy tracking SynPAR$_{TF}$ localization with and without activation. (A)** TEV$_{cs}$ and Null$_{cs}$ Chemo-SynPAR$_{TF}$ were transiently transfected into HEK 293T cells with NanoLuc-arrestin-TEVp and UAS-reporter. Cells were primed with TEV$_p$ and stimulated with SalB and light prior to fixation, permeabilization, and staining. The Null$_{cs}$ construct is membrane localized after activation while the TEV$_{cs}$ construct in internalized. **(B)** TEV$_{cs}$-AutoSynPAR$_{TF}$ also shows activation-dependent internalization. Untreated receptor (top) is predominantly localized to the cell surface while TEV$_p$ treated receptor (bottom) is predominantly internalized. Representative fields of view imaged at 63X. Scale bars are 10 microns.

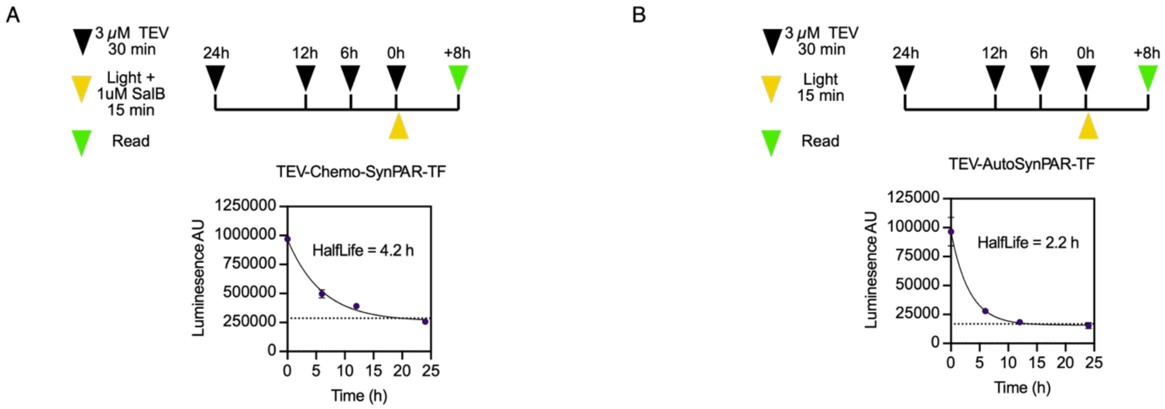


**Supplementary Figure 4 Decay of SynPAR$_{TF}$ signal with variable time between prime and stimulation steps. (A)** Cells expressing TEV$_{cs}$-Chemo-SynPAR$_{TF}$ were primed with TEV$_p$ for 30 min at different timepoints (black arrows) prior to stimulation with light and SalB (yellow arrows). Reporter expression was read out 8h after stimulation (green arrows). **(B)** Cells expressing TEV$_{cs}$-Auto-SynPAR$_{TF}$ were primed with TEV$_p$ for 30 min at different timepoints (black arrows) prior to stimulation with light (yellow arrows). Reporter expression was read out 8h after stimulation (green arrows). Data plotted as mean of triplicate, with error plotted as SD. This experiment was performed once.

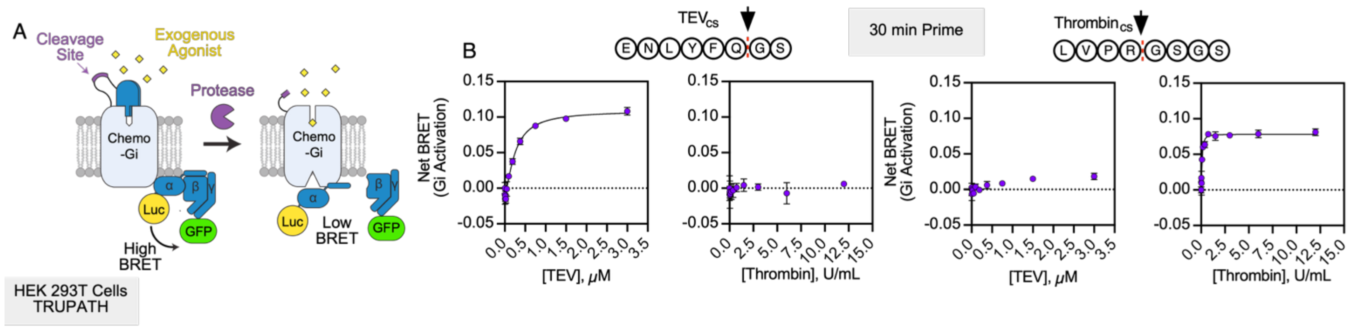


**Supplementary Figure 5** Chemo-SynPAR$_{Gi}$ responds to programmed protease inputs. **(A)** A schematic of Chemo-SynPAR$_{Gi}$, which can couple to Gi signaling as detected by the TRUPATH BRET-based assay. **(B)** Protease dose-response experiments demonstrate that Chemo-SynPAR$_{Gi}$ can respond to priming with either TEV or thrombin in a dose-dependent manner when subsequently stimulated by 1 nM DCZ chemical agonist.

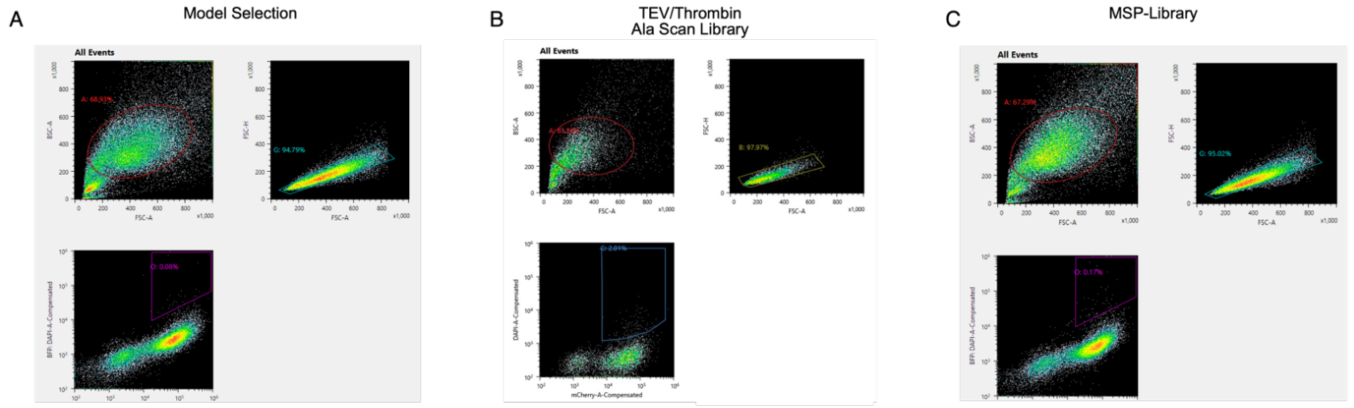


**Supplementary Figure 6** **(A-C)** Gating strategies for model selection **(A)**, TEV/Thrombin alanine scan **(B)**, and MSP-Library sorts **(C)**. Top plots are scattering gates for viability and single cells. For bottom plot, Y axis is tBFP expression, X axis is mCherry expression. All samples were stimulated with SalB and Light but not protease (i.e. background).

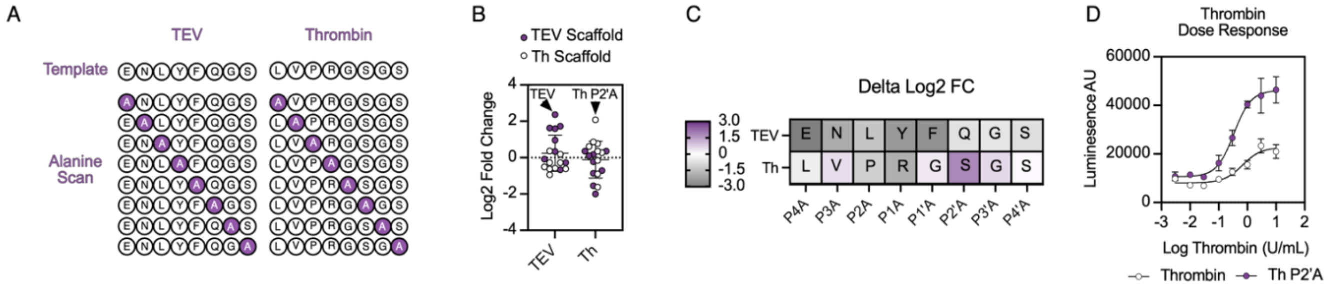


**<u>Supplementary Figure 7</u> Library-based selections with Chemo-SynPAR$_{TF}$ can capture determinants of protease activity and enhance SynPAR function. (A)** A 19 member library was designed and inserted into the Chemo-SynPAR$_{TF}$ scaffold for pooled screening in the TARGATT 293-based selection system. **(B)** Analysis of enrichment of TEV$_{cs}$ (purple) and Thrombin$_{cs}$ (white) scaffolds for TEV (left) and thrombin (Th, right) treated samples. **(C)** Analysis of the relative change in enrichment for a given alanine mutant relative to the template scaffold. Higher fold change indicates that Alanine is preferred at that position **(D)** A comparison of Thrombin$_{cs}$ and its P2'A mutant in a transient luciferase reporter assay after priming with different thrombin concentrations and stimulation with light and 1 uM SalB.

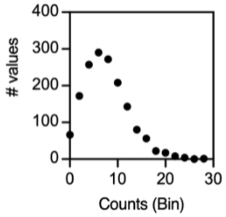


**<u>Supplementary Figure 8</u>** A frequency distribution for the MSP input library comparing counts (x axis) and representation of sequences with that level of abundance (y axis). Most library members were present at 5-10 copies in the MSP library.

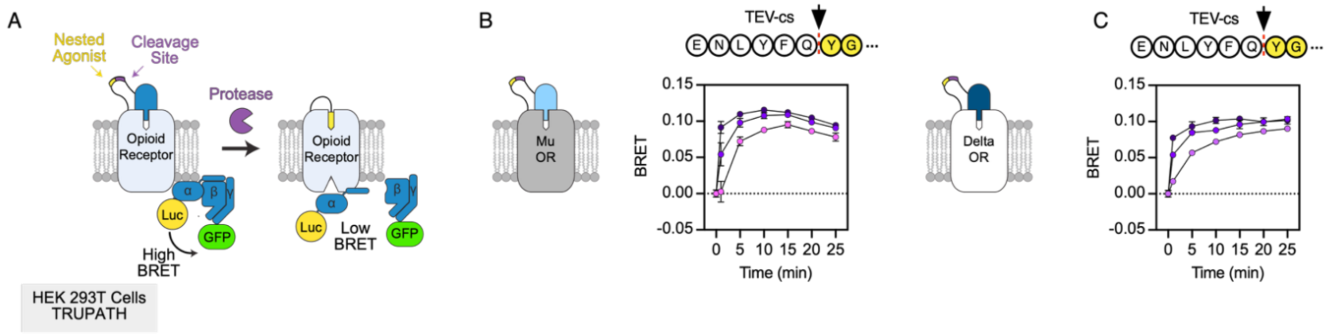


**Supplementary Figure 9** **(A)** Schematic of Auto-SynPAR$_{Gi}$, which can couple to Gi signaling as detected by the TRUPATH BRET-based assay. **(B-C)** TEV$_{cs}$-Auto-SynPAR$_{Gi}$ construct based on μOR **(B)** and δOR **(C)** generate saturable, time- and dose-dependent activation of Gi in HEK293T cells as detected by the TRUPATH BRET-based assay. Data plotted as mean of triplicate, with error plotted as SD.

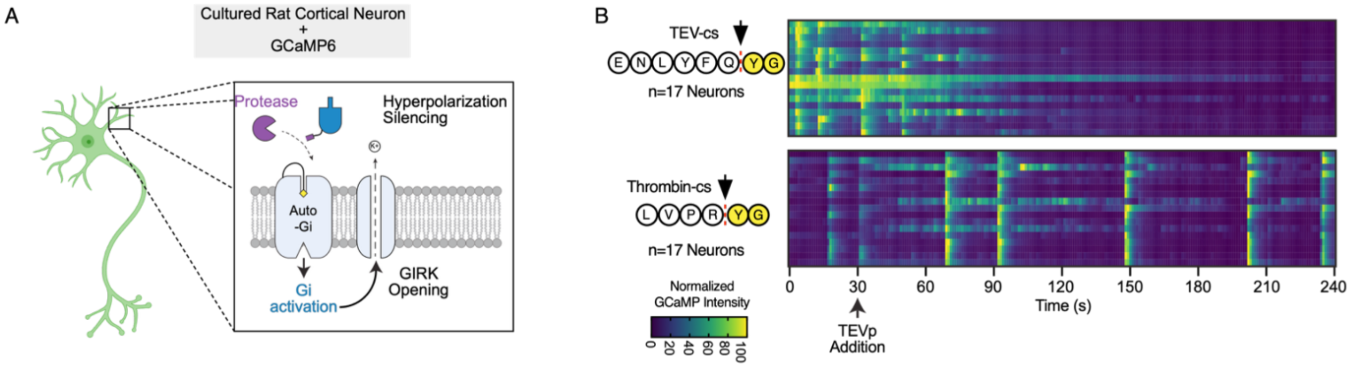


**Supplementary Figure 10** **(A)** Schematic of a cultured rat cortical neuron expressing the calcium sensor GCaMP6 and a κOR-based Auto-SynPAR-Gi construct to facilitate protease specific silencing by activating GIRK channels in response to Tryptic protease activity. **(B)** Plots of normalized GCaMP intensities of rat cortical neurons (n=17) expressing $PAR2_{cs}$-Auto-$SynPAR_{Gi}$ (top) or $TEV_{cs}$-Auto-$SynPAR_{Gi}$ (bottom). Treatment of neurons with TEV protease (3 uM at t=30s, arrow) at DIV 14 was sufficient to specifically silence spontaneous firing in neurons expressing the matched SynPAR construct. All experiments are representative of at least biological duplicates.

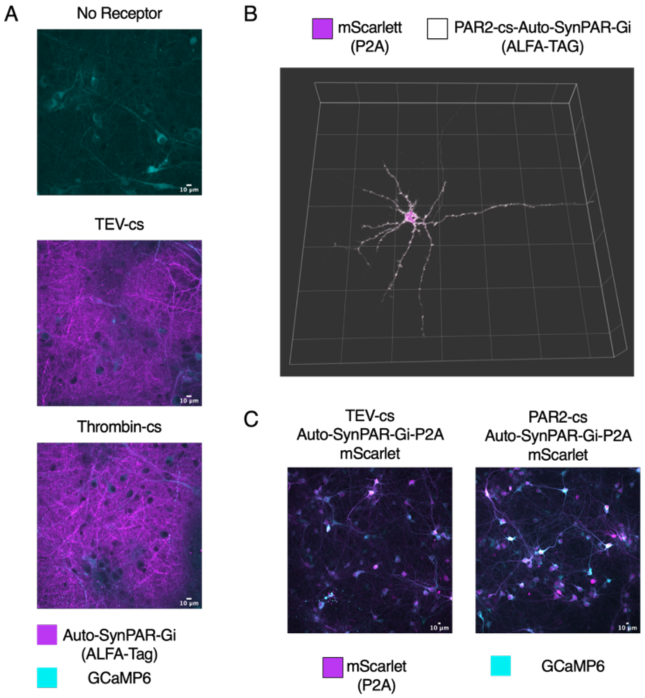


**Supplementary Figure 11** **Assessment of Auto-SynPAR$_{Gi}$ constructs in cultured rat cortical neurons. (A)** Staining of neurons at 14 DIV that were transduced with AAV1/2 at 7 DIV containing hSyn_Auto-SynPAR$_{Gi}$ (Visualized with Anti-ALFA-Tag-647, magenta) and hSyn_GCaMP6 (cyan) constructs as indicated. **(B)** A z-stack reconstruction of a single neuron at 14 DIV that was transfected with hSyn_PAR2$_{cs}$-Auto-SynPAR$_{Gi}$-P2A mScarlet (magenta) and stained with Anti-Alfa-Tag-647 (white). **(C)** Visualization of cultured rat cortical neurons at 14 DIV that were transduced with AAV1/2 at 7 DIV containing hSyn_Auto-SynPAR$_{Gi}$ -P2A mScarlet (magenta) and GCaMP6 (cyan). All scale bars are 10 microns.

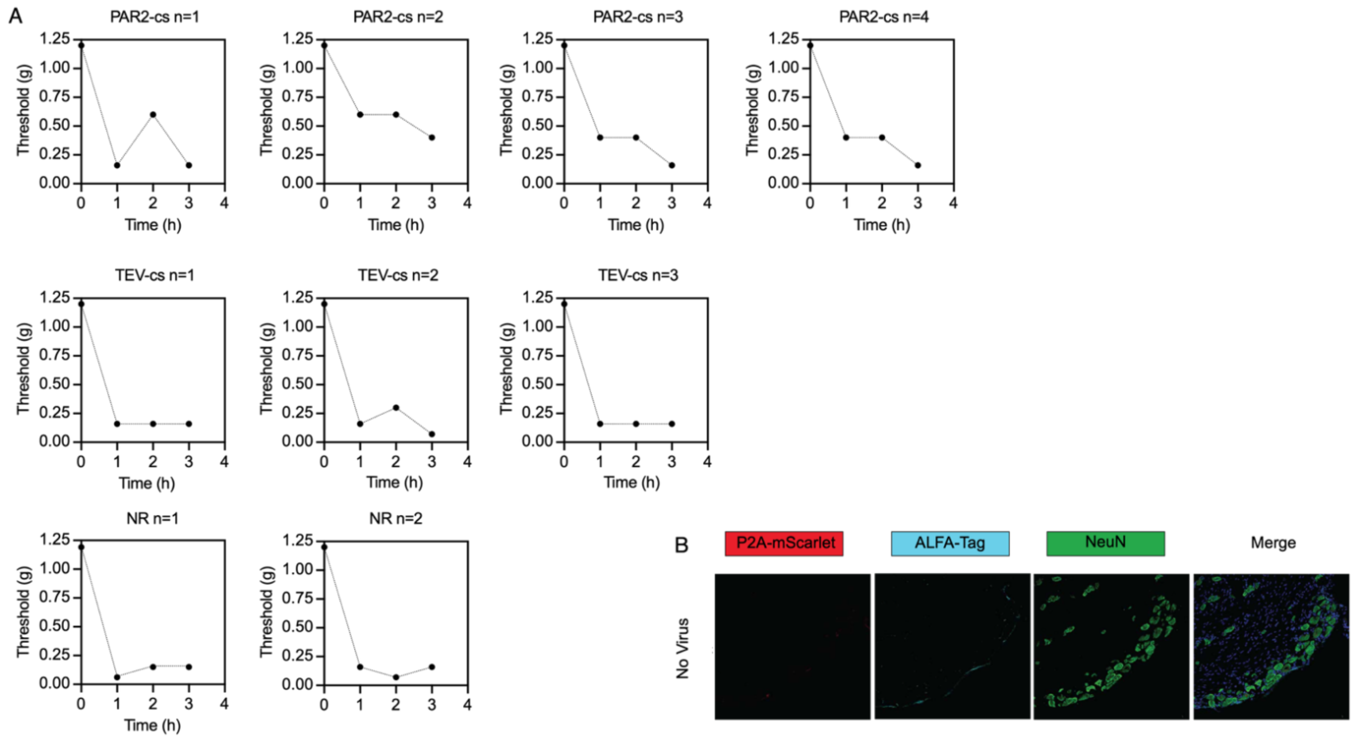


**<u>Supplementary Figure 12</u> (A)** Individual mouse data for VFF test for mechanical hyperalgesia in Figures 5D and E. **(B)** DRG staining for representative animal with that received a sham injection rather than virus (NR, no receptor control).
